# Cross-Fostering with control dams rescues Gut Dysbiosis and Chromatin-associated Transcriptional Changes in Offspring of Opioid-Exposed Dams

**DOI:** 10.1101/2025.11.07.687278

**Authors:** Somnath Pandey, Yaa F. Abu, Praveen Singh, Sabita Roy

## Abstract

Prenatal opioid exposure disrupts gut homeostasis and causes gastrointestinal complications in offspring, but the mechanisms remain unclear. Here using a murine model of prenatal hydromorphone exposure, we examined gut microbiota, intestinal injury, transcriptomic signatures, and chromatin accessibility. Exposed pups displayed marked dysbiosis, epithelial damage, and upregulation of inflammatory gene programs accompanied by relaxed ileal chromatin. Cross-fostering to opioid-naïve dams restored microbial diversity, reestablished metabolite-producing taxa, and reversed injury-associated transcriptional and chromatin changes. Fecal microbiota transplantation from exposed dams recapitulated intestinal injury, indicating a microbiome-driven mechanism. These findings reveal a novel gut-microbiome-epigenome axis underlying opioid-induced injury and highlight early microbial intervention as a potential strategy to mitigate developmental harm.

## Introduction

The rise in infants with prenatal histories of opioid exposure is an indirect consequence of the opioid epidemic [1]. Recent reports indicate that nearly 7% of women use prescription opioids during pregnancy. Of those, approximately 20% report misuse, defined as obtaining opioids outside medical care or using them for non pain purposes [2]. The number of pregnant women with opioid-related diagnoses increased from 3.5 to 8.2 per 1000 delivery hospitalizations between 2010 to 2017 [3]. Neonatal opioid withdrawal syndrome (NOWS) is a well-described consequence of sudden opioid withdrawal in offspring with symptoms affecting the central and autonomic nervous systems, gastrointestinal tract, and respiratory system [1, 2, 4–6]. Importantly, the effects of prenatal opioid exposure extend beyond the neonatal period, increasing the risk for developmental, behavioral, educational, and mental health challenges later in life [4].

It is increasingly evident that adverse outcomes in opioid-exposed offspring result from a complex interplay of biological and environmental factors. The transmission of risk is intergenerational, encompassing genetic and epigenetic changes as well as social, psychological, and physical stressors encountered during childhood [5–7]. For this reason, animal models are a convenient means of examining effects of prenatal opioid exposure itself without the confounds present in heterogeneous clinical populations. Indeed, animal models of prenatal opioid exposure are slowly accumulating, and have described biological consequences of *in-utero* opioid exposure, which parallel many findings in clinical models [7–9]. Additionally, the preclinical field is slowly advancing towards an appreciation of mechanisms underlying these findings. For instance, Jantzie et al., [10, 11] reported peripheral inflammation, immune priming, and sustained peripheral immune reactivity in opioid-exposed offspring. Our laboratory has established a connection between prolonged morphine use and microbial dysbiosis in adult mouse models [12–14]. Furthermore, the lab has linked this microbial dysbiosis to systemic inflammation [14–16]. These findings are particularly relevant for women of reproductive age who are using opioids, as they may develop and sustain microbial dysbiosis that persists throughout the course of pregnancy. While it has not yet been directly demonstrated, there is growing support for the possibility that prenatal exposure to opioids may result in gut microbial dysbiosis in offspring.

Several lines of evidence underpin this hypothesis-1) there is a well-documented, strong association between the mother’s gut microbial composition and diversity and that of her infant [17]; 2) NOWS, which often occurs following in utero opioid exposure, is characterized by gastrointestinal disturbances and neurological symptoms observable at birth, likely mediated through the gut-brain axis [18–20]; 3) Research has identified the presence of a gut microbiome during the prenatal period [21, 22]. Opioid exposure during pregnancy may disrupt this early microbiome, potentially leading to alterations in immune system development. This is particularly significant because immune development is closely intertwined with the early establishment and maturation of the gut microbiota [23]. Although the detrimental consequences of maternal opioid use on offspring—manifesting as NOWS—are well-documented, the underlying etiological mechanisms responsible for these adverse outcomes remain incompletely understood. Emerging clinical and translational research suggests that both heritable genetic predispositions and environmentally responsive epigenetic modifications play pivotal roles in modulating the severity and variability of NOWS phenotypes. In particular, prenatal exposure to opioids is hypothesized to influence epigenetic programming, thereby altering gene expression profiles critical to gut-development and stress regulation [24].

Recent studies, including our own, have demonstrated that the gut microbiome plays a crucial role in mediating the adverse effects of prenatal opioid exposure [10–13], encompassing immunological, gastrointestinal, and other detrimental outcomes in offspring [21, 25]. Opioid use during pregnancy causes gut disruption and similarly impacts the microbiome in dams and offspring, with lasting changes in gut microbial composition [21, 26]. The gut microbiome interacts with the host transcriptome and epigenome, influencing various outcomes, particularly appreciated in inflammatory diseases [27, 28]. While changes in gut microbiota diversity following prenatal opioid exposure are documented, little is known about how it influences the intestinal homeostasis and the global chromatin landscape.

In this study, we use a murine model of prenatal hydromorphone exposure and cross-fostering to dissect the relationships among the gut microbiome, host transcriptome, and directly assessing the chromatin accessibility landscape. We also distinguished prenatal versus postnatal effects through cross fostering and effects on the microbiome. Cross-fostering after birth represents a window of therapeutic intervention that can be leveraged to induce a lasting shift in the neonatal gut microbiome and associated health outcomes.

## Methods

### Animals

All animal experiments were approved by the Institutional Animal Care and Use Committee policies at the University of Miami and adhered to all ethical guidelines related to the care of laboratory animals. Twelve-week-old female C57 BL/6 mice (25-30g) were purchased from Jackson Laboratories (Bar Harbor, ME, USA) (strain 3752), with age-matched littermate wild type mice used as controls. Mice were housed five per cage under a controlled temperature (22 ± 2C), humidity (30%–70%), and 12 h light/dark cycle (light at 0700), with food pellets and water ad libitum. Mice were maintained in sterile microisolator cages under pathogen-free conditions. At the conclusion of experiments, mice were humanely sacrificed using CO2 asphyxiation followed by cervical dislocation, as recommended by the Panel of Euthanasia of the American Veterinary Medical Association (AVMA).

### Prenatal Opioid Exposure

iPRECIO 310R infusible mini pumps (SMP-310R, Primetech, Tokyo, Japan) filled with hydromorphone (prenatally opioid-exposed group) or saline (control group) were subcutaneously implanted into the nape of 12-week-old female C57BL/6 mice. Programming was set to continuously deliver hydromorphone or saline at a flow rate of 1ul/hour. Hydromorphone (or saline) was refilled in pumps every five days at starting dose of 100ug/day, with subsequent doses increasing by 50 ug/week for hydromorphone treatment. Mice were mated for 7 days with drug naive males until confirmation of pregnancy, assessed by presence of a copulation plug. Hydromorphone treatment through infusible mini pumps continued until pups were born on postnatal day 1 (P1), after which drug release through pumps was immediately discontinued. This model intends to recapitulate pre-gestational initiation of opioids followed by gestational exposure alone to opioids, with no postnatal opioid exposures in offspring. In some experiments, P1 pups stayed with their birth mothers—hydromorphone-exposed pups nursed by hydromorphone mothers (HP_HM) or control pups nursed by control mothers (CP_CM) —and were weaned on P21. In others, some P3 pups were cross-fostered to lactating dams —hydromorphone-exposed pups nursed by control mothers (HP_CM) or control pups nursed by hydromorphone mothers (CP_HM) and weaned on P21 for experiments. Experiments were conducted in male and female offspring but were not powered to detect sex-differences.

### 16S rRNA gene sequencing

DNA was extracted from small intestinal contents using DNeasy PowerSoil Pro Kit (Qiagen, Maryland, USA). Sequencing was performed by the University of Minnesota Genomics Center. The V4 region of the bacterial 16S rRNA gene was amplified with primers 515F/806R and sequenced on an Illumina MiSeq (2 × 250 bp). Raw reads were processed in R (v4.x) using the DADA2 pipeline: primers were trimmed, reads were quality-filtered (maxEE = 2), denoised, merged, and chimeras removed to generate amplicon sequence variants (ASVs). Taxonomy was assigned against the SILVA v138 reference database. A phylogenetic tree was inferred under the GTR+G model using IQ-TREE (v2.x). BugBase is a microbiome analysis algorithm that predicts high-level phenotypes present in microbiome samples using 16S amplicon data. The BugBase phenotype predictions were implemented using the online web app (https://bugbase.cs.umn.edu/).

### Microbiome data integration and filtering

The ASV abundance table, taxonomy, sample metadata and phylogeny were combined into a phyloseq object. Samples with total reads < 1,000 and taxa lacking phylum-level assignment were excluded. Taxa present in < 5% of samples were filtered out. For compositional analyses, ASV counts were converted to relative abundances.

### PICRUST Functional Prediction

Representative ASV sequences and an ASV abundance table were exported from DADA2 (seqtab.nochim.Rds) and formatted for PICRUSt2. ASV sequences were written to rep_seqs.fasta and the OTU table to OTUdf_clean.tsv (ASV IDs as row names, no quotes). PICRUSt2 v2.4.1 was run via the picrust2_pipeline.py wrapper (8 threads), which performs: (1) Sequence placement (place_seqs.py) of ASVs into a reference phylogeny, (2) Hidden-state prediction (hsp.py) of gene family copy numbers for each ASV, corrected by 16S rRNA gene copy number, (3) Metagenome inference (metagenome_pipeline.py) to predict per– sample KEGG Ortholog (KO) abundances, and (4) Pathway inference (pathway_pipeline.py) using MinPath against the MetaCyc database to generate predicted pathway abundances. The resulting per-sample pathway abundance table was imported into R, converted to relative abundances, and compared between CP_CM (n = 8) and HP_HM (n = 8) groups or HP_CM (n = 8) and HP_HM (n = 8). Two-sided Welch’s *t*-tests were performed for each pathway, with Benjamini–Hochberg correction for multiple comparisons. Pathways with FDR-adjusted *p* < 0.05 were considered significantly different.

### Diversity analyses

Alpha diversity metrics (Observed richness, Shannon, Simpson, Chao1) were computed on rarefied data (rarefy_even_depth, seed = 123) using *estimate_richness*. Group comparisons of alpha diversity were performed by two-sided Student’s *t*-test. Beta diversity was calculated by Bray–Curtis dissimilarity, visualized via principal coordinates analysis (PCoA) with 95% confidence ellipses, and tested by PERMANOVA (*adonis2*, 999 permutations).

### Differential abundance testing

At phylum, family, genus and species levels, relative-abundance data were tested for pairwise differences between CP_CM and HP_HM or HP_CM and HP_HM, using two-sided *t*-tests with Bonferroni correction (adjusted *p* < 0.05). Significant taxa were visualized as boxplots.

### ATAC-sequencing

Small intestinal tissues were processed, nuclei isolated, followed by tagmentation, and sequenced by Active Motif Inc., using the manufacturer’s recommendations (Cat. No. 53150). ATAC-sequencing pipeline includes quality control and adapter trimming, alignment to a reference genome, peak calling, peak annotation, differential accessibility analysis, and visualization of results. The quality control and adapter trimming steps ensure that low-quality and adapter sequences are removed from the raw data. Alignment to the reference genome is performed using the Bowtie2 algorithm, which allows for the detection of open chromatin regions in the genome. Peak calling is performed using MACS 2.1.0 [PMID: 18798982]. For the comparative analysis standard normalization is achieved by down sampling the usable number of tags for each sample in a group to the level of the sample in the group with the fewest usable number of tags. Differential accessibility analysis is performed using DESeq2 on raw peak counts, with peaks considered significant at unadjusted *p*<0.05 and |shrunken log₂ fold change|>0.5, which allows for the identification of differentially accessible regions between two or more conditions.

### RNA-seq

RNA from experiments in biological triplicates was isolated using RNeasy Mini Plus Kit (QIAGEN) following the manufacturer’s instructions. Strand-specific RNA libraries were generated from 1 μg of RNA using TruSeq stranded total RNA with Ribo-Zero Gold (Illumina). Sequencing was performed using single end reads (150 bp, average 50 million reads per sample) on the HiSeq2500 platform (Illumina) at Active Motif. Sequenced reads were aligned to the mm10 genome assembly using TopHat2 (<u>Kim et al., 2013</u>). The expression level and fold change of each treatment group was evaluated using DESeq2 [PMID: 25516281]. Genes that had 0 reads across all samples were excluded. In order to get rid of batch effects, samples were normalized using RUVr method from the RUVseq package (<u>Risso et al., 2014</u>). For the repetitive element analysis, RNAseq reads were mapped against known repetitive element sequences and data were normalized with the total number of mappable reads. Significantly regulated genes were defined as log2FC >1 or < -1, and FDR < 0.5.

### Pathway enrichment

We used the *p*<0.05, |LFC|>1.0 gene list (n = X genes) for Ingenuity Pathway Analysis (IPA, Qiagen). Canonical pathway enrichment *p*-values (Fisher’s exact test) and activation *z*-scores were calculated; the top pathways by –log₁₀(*p*) are presented, with *z*-score direction indicating predicted activation (>0) or inhibition (<0).

### Fecal Microbial Transfer (FMT)

Fecal contents from saline treated control dams or hydromorphone exposed dams mice were collected after sacrifice. The fecal content was processed, as previously described [21]. Briefly, 200 mg of fecal extracts were suspended in 1 mL sterile anaerobic solution, vortexed to homogenize, and filtered through 70 µM cell strainer, and finally, centrifuged briefly for 2 min. 3-week old naïve C57BL/6 mice were randomized into 2 treatment groups (hydromorphone plus saline treatment group and hydromorphone only group) This was followed by oral gavage with 200 µL of freshly prepared fecal suspension for 7 consecutive days. Mice were transferred to clean cages after FMT given mice coprophagic behavior. On day 8, the mice were humanely euthanized and ileal tissue was harvested. Tissues were subjected to H&E staining

### Data and Statistical Analysis

Statistical analysis and figures were generated using R software (v4.x) with the packages *phyloseq*, *vegan*, *ggplot2*, *ggpubr*, *microbiome*, *microViz*, *DESeq2*, *rstatix*, *ComplexHeatmap* and *patchwork*.

## Results

### Prenatal hydromorphone exposure significantly disrupts gut microbial composition with concomitant reduction in bacterial community implicated in maintaining gut health

Opioid exposure has previously been shown by our laboratory to induce microbial dysbiosis and intestinal injury in adult mice [29]. To investigate the effects of prenatal hydromorphone exposure on the gut microbiome, stool samples from 3-week-old offspring prenatally exposed to hydromorphone and nursed by hydromorphone mothers (HP_HM) or from control offspring nursed by control mothers (CP_CM) (Fig 1A) were subjected to 16S rRNA sequencing. Alpha diversity per Shannon index was significantly decreased with prenatal hydromorphone exposure (Fig 1B). Similarly, distinct clustering between CP_CM and HP_HM fecal samples were visualized on a principal component analysis (PCoA) plot, suggesting significant differences in β-diversity per the Bray-Curtis dissimilarity index (PERMANOVA: F = 5.595, R2 = 0.359, *p* = 0.004) (Fig 1C). At the phylum level, prenatal hydromorphone exposure significantly lowered the mean relative abundance of *Actinobacteria* (p =0.049) and *Bacteroidota* (p=0.046), while significantly increasing the relative abundance of *Firmicutes* (p=0.002) (Fig 1D). Differential abundance analysis at the genus level revealed reduction in 6 genera (Fig 1E-J). The relative abundance of *Bifidobacterium, Romboutsia,* and *Lachnospiracea A2* significantly decreased in the HP_HM group compared to CP_CM. Conversely, the relative abundance of genera *[Eubacterium] xylanophilium group*, *Lachnospiraceae UCG-006*, and *Lachnoclostridium* were enriched in HP_HM relative to CP_CM. To understand the functional capabilities of microbial communities in the gut microbiome and predict the phenotypic traits of microbes, we probed our 16S rRNA sequencing data. We utilized Bugbase algorithm to investigate the changes observed in the high-level phenotypes in the microbiome. The analysis revealed that prenatal hydromorphone exposure is linked to changes in bacterial composition corresponding to an increase in the predicted relative abundance of facultative anaerobic, gram positive, and mobile-element-containing bacteria, and a decrease in the predictive relative abundance of gram negative and biofilm-forming bacteria (Supplementary Table 1A). To explore the functional profile of the microbial community based on 16S rRNA sequencing, we resorted to PICRUST pathway analysis. Changes in bacterial taxa were associated with significant differences in the metabolic profile of over 30 distinct pathways per PICRUST analysis (Supplementary Figure 1). To note, pathways relevant for Biotin Biosynthesis and for Short-chain Fatty acid synthesis (SFCAs) such as the Gamma-Aminobutyric Acid Shunt, Lactose and Galactose Degradation Pathway, Anhydromuropeptides Recycling were found to be significantly reduced following prenatal hydromorphone exposure (Fig 1K). Additionally, the PICRUST analysis revealed changes in pathways relating to pyruvate fermentation to Butanoic acid, L-methionine biosynthesis, the TCA cycle, tRNA processing, thiamin salvage, pyrimidine deoxyribonucleotides de novo biosynthesis, the NAD salvage pathway were predicted to be significantly downregulated with prenatal hydromorphone exposure (Supplementary Figure 1). These pathways collectively underscore the relevance of microbial metabolic functions in establishing prenatal microbiomes in mice. They contribute to nutrient synthesis (e.g., biotin, etc.), energy production and gut maintenance (e.g., SCFAs, TCA cycle, etc.), immune modulation, neurodevelopmental support (e.g., GABA metabolism), and microbial diversity maintenance. Whereas pathways related to glycolysis V and the NAD salvage pathway II which are mostly associated with stress response, were predicted to be significantly upregulated.

**Figure 1.**
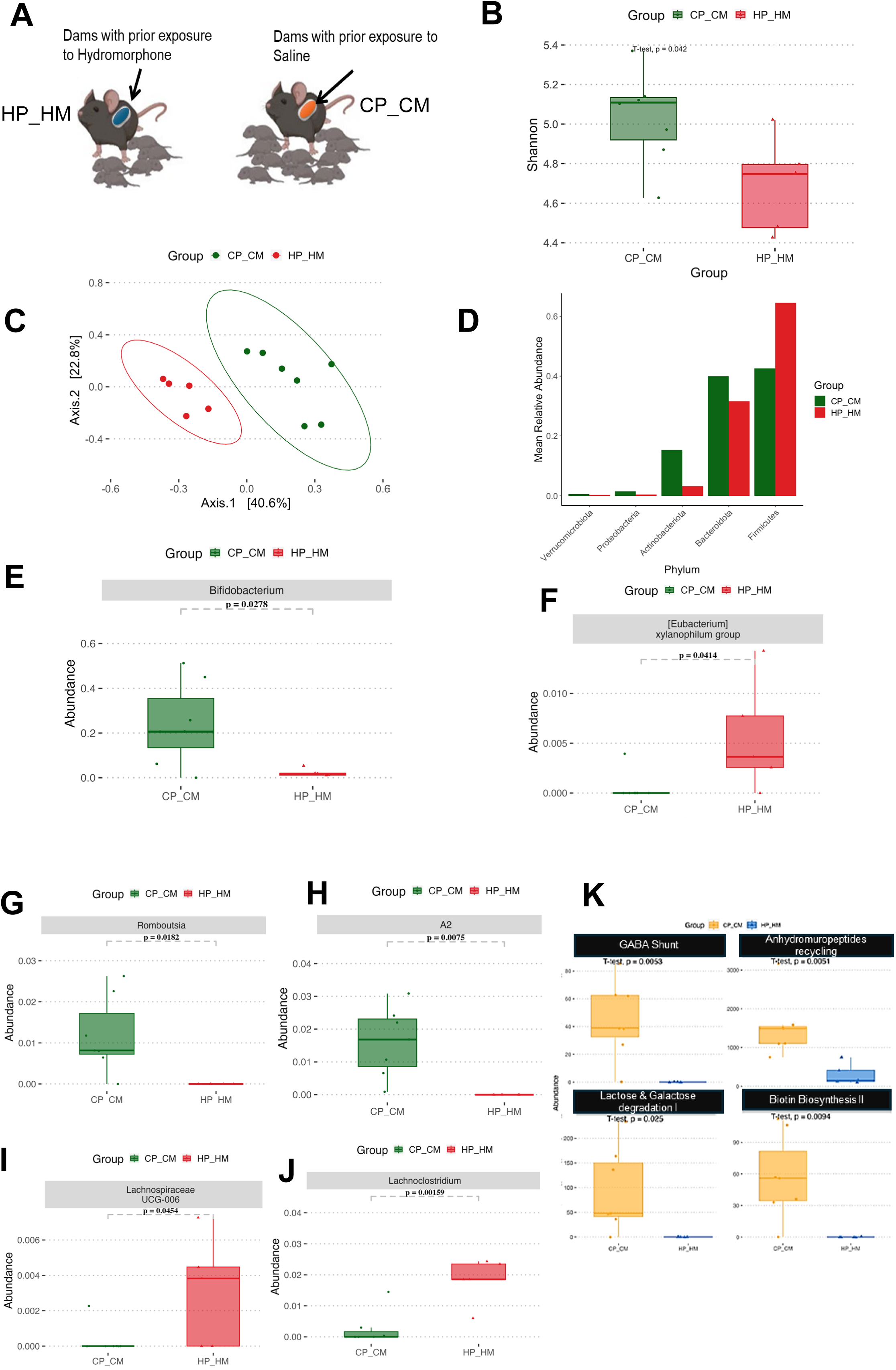
Prenatal hydromorphone exposure significantly disrupts gut microbial composition with concomitant reduction in bacteria implicated in maintaining gut health. **(A)** Generation of prenatal opioid-exposed (HP_HM) or control (CP_CM) offspring from opioid exposed (hydromorphone mom) or saline exposed (control mom) dams. **(B)** Shannon diversity index for CP_CM (green) and HP_HM (red) groups; boxes show interquartile range, whiskers 1.5 × IQR, and center lines the median; *p* < 0.05 by two-sided *t*-test. **(C)** Principal coordinates analysis (PCoA) of Bray–Curtis distances; each point is a sample and ellipses denote 95% confidence intervals, demonstrating clear separation of CP_CM and HP_HM microbiota. **(D)** Bar plots of relative abundances of major bacterial phyla in each sample, grouped by CP_CM and HP_HM; only phyla with mean abundance > 1% are shown. **(E–J)** Boxplots of select genera with significant differential abundance between groups: **(D)** *Bifidobacterium*, **(E)** *Romboutsia*, **(F)** A2, **(G)** *[Eubacterium] xylanophilum* group, **(H)** *Lachnospiraceae_UCG-006*, **(I)** *Lachnoclostridium*. Boxes and whiskers as in (A); adjusted *p* < 0.05 by Bonferroni-corrected *t*-test. **(K)** PICRUST functional analysis. Boxplots showing relative abundance of four representative pathways; *p*-values by Bonferroni-adjusted pairwise *t*-tests.

In total, these data provide evidence of dysbiosis associated with prenatal hydromorphone exposure and the reduced presence of good bacteria implicated in the synthesis of SCFAs and other such metabolites, implicated in maintaining gut health.

### Prenatal hydromorphone exposure is associated with skewed transcription and chromatin profiles corresponding to intestinal tissue damage and reduced homeostatic-stress mechanisms

The gut microbiome plays a role in maintaining intestinal homeostasis, in part through transcriptional and epigenetic regulation of the intestinal epithelium [30]. Given that prenatally hydromorphone exposed offspring displayed microbial dysbiosis, we next investigated the impact of hydromorphone exposure on small-intestinal gene expression. To determine whether tissue-specific transcriptional signature exists with prenatal hydromorphone on a global scale, bulk RNA-sequencing on distal-ileal tissue was performed. Principal component analysis was used to illustrate differential clustering between HP_HM and CP_CM groups (Fig. 2A). Differential gene expression analysis revealed several genes significantly upregulated or downregulated with prenatal hydromorphone exposure in the volcano plot (Fig. 2B). In particular, 686 genes were upregulated and 765 genes were downregulated in the HP_HM group relative to CP_CM (adjusted p-value <0.05 & |log2FC| >0.5).

**Figure 2.**
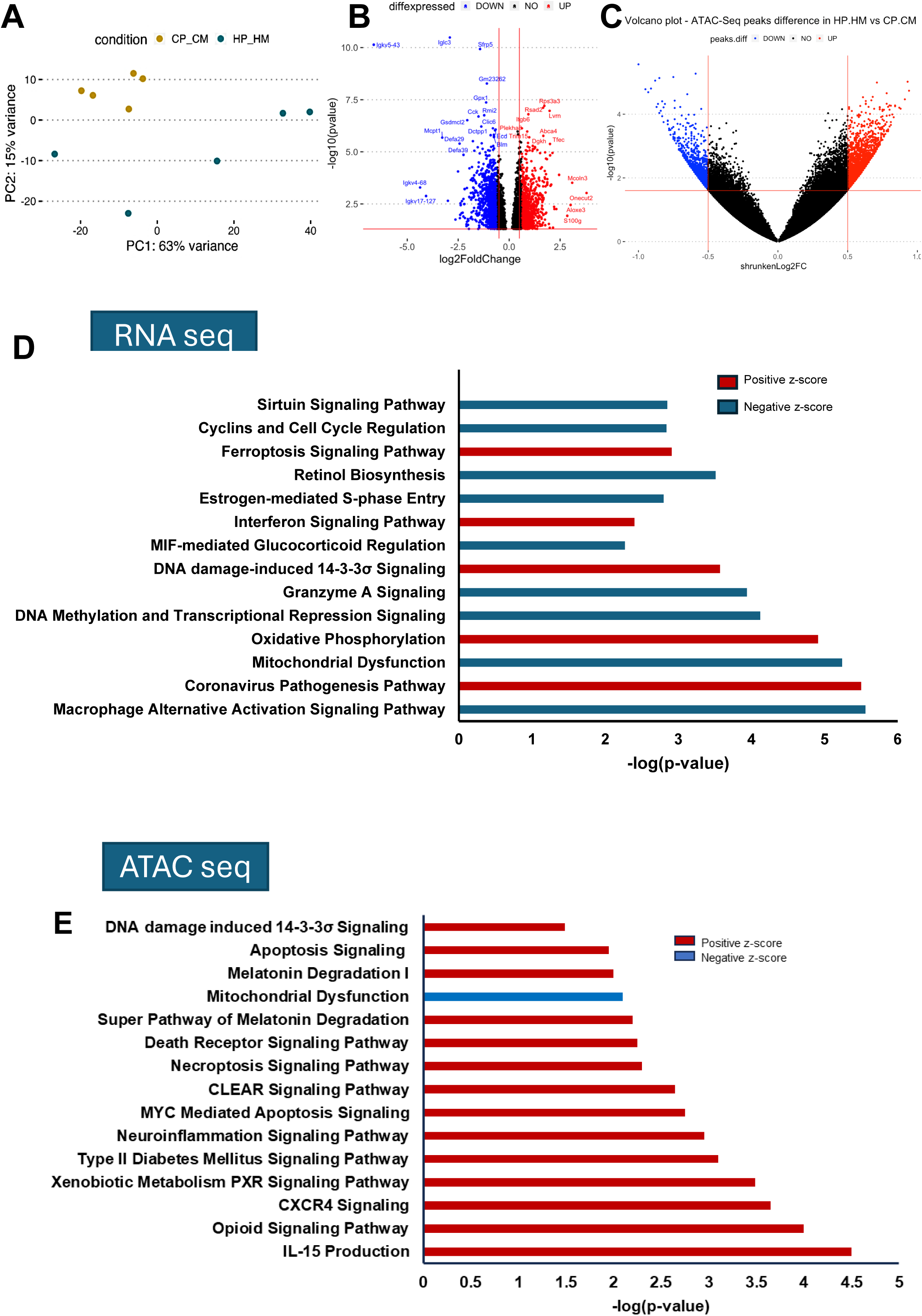
Prenatal hydromorphone exposure shows skewed transcription and chromatin profiles corresponding to intestinal tissue damage and reduced homeostatic-stress mechanisms (n=5/group) **(A)** PCA of variance-stabilized RNA-Seq counts separates CP_CM (gold) and HP_HM (teal) samples along PC1 (63% variance) and PC2 (15%). **(B)** Volcano plot of differential gene expression (HP_HM vs CP_CM): log₂ fold change vs – log₁₀(*p*). Genes with *p*<0.05 and |LFC|>0.5 are colored red (up in HP_HM) or blue (down); selected DEGs are labeled. **(C)** Volcano plot of differential ATAC-Seq peak accessibility (HP_HM vs CP_CM): shrunken log₂ fold change vs –log₁₀(*p*). Peaks with *p*<0.05 and |LFC|>=0.5 are highlighted in red (more accessible in HP_HM) or blue (less accessible). **(D)** Top enriched canonical pathways from IPA based (from RNA-seq between HP_HM vs CP_CM) on the unadjusted *p*<0.05, |LFC|>1.0 gene set. Bars denote –log₁₀(*p*); red, positive *z*-score (predicted activation in HP_HM); blue, negative *z*-score (predicted inhibition). Because FDR filtering returned too few genes for reliable pathway calls, unadjusted *p* was used to expand the gene set. **(E)** Top enriched canonical pathways from IPA based on the unadjusted *p*<0.05, of genes with differential ATAC peaks in HP_HM versus CP_CM. Bars denote –log₁₀(*p*); red, positive *z*-score (predicted activation in HP_HM); blue, negative *z*-score (predicted inhibition). Because FDR filtering returned too few genes for reliable pathway calls, unadjusted *p* was used to expand the gene set.

To elucidate the biological pathways and molecular mechanisms underlying the effects of prenatal hydromorphone exposure, Ingenuity Pathway Analysis (IPA) was performed on the differentially expressed gene set. For pathways discovery, our IPA analysis on the unadjusted *p*<0.05, |LFC|>1.0 gene set, showed upregulation of the DNA damage-induced 14-3-3σ Signaling pathway, oxidative phosphorylation signaling, Ferroptosis pathway, Interferon Signaling, and Coronavirus pathogenesis pathway in the HP_HM group compared to the CP_CM (figure 2D). Notably, increased oxidative phosphorylation signaling has been shown to occur with tissue damage [31–34]. On the other hand, pathways such as the macrophage alternative activation signaling pathway—which plays an instrumental role in orchestrating immune regulation, wound healing, and repair [35, 36] — was downregulated in the HP_HM group, indicating lack of repair mechanisms or excessive damage in small intestinal tissue when compared to the CP_CM group. Similarly, pathways affecting Retinol biosynthesis, Mitochondrial function, Granzyme A signaling, MIF-mediated Glucocorticoid regulation, Cell cycle regulation, estrogen-mediated S-phase entry, and the homeostatic-stress responsive, Sirtuin signaling pathway, were found to be downregulated in the HP_HM group (figure 2D). Earlier observations have shown that mitochondrial abnormalities/diseases are repercussions of fundamental defect in oxidative phosphorylation [37, 38]. Our RNA-seq analysis therefore revealed pathways enriched in inflammation, tissue damage and reduced homeostatic-stress response in the HP_HM group relative to CP_CM group.

We hypothesized that changes in gene transcription correlating to distal intestinal tissue inflammation and damage noted above are caused due to alterations in chromatin accessibility of associated genes in prenatal hydromorphone exposure group. To understand the potential cause of altered transcription levels between sample groups, the chromatin architecture status of these samples was assessed using the Assay for Transposase-Accessible Chromatin (ATAC-seq) sequencing, to gain insight on genome-wide chromatin accessibility. We applied the Ingenuity Pathway Analysis platform on the accessible chromatin regions identified through ATAC-seq. To ensure the robustness of our ATAC-seq analysis, we first probed the chromatin accessibility status of a House Keeping Gene such as beta-Actin (Supplementary Fig 2). As expected, we observed that the peaks for HP_HM versus CP_CM groups were the same and not impacted by the treatment at this control gene loci. Further, our analysis revealed increased accessibility to genes corresponding to pathways implicated in inflammation such as IL-15 production, CXCR4 signaling, neuroinflammation signaling, and Diabetes Type II signaling pathway in the HP_HM group (figure 2E). Satisfyingly, we observed that prenatal hydromorphone exposure is associated with alterations in the chromatin architecture, more specifically, increased accessibility to genes involved in Opioid signaling and Xenobiotic metabolism PXR Signaling in the HP_HM group. Additionally, genes implicated in pathways involved in cell death such as MYC mediated Apoptosis Signaling, Necroptosis Signaling pathway, Death Receptor Signaling, and Apoptosis Signaling showed more accessibility in HP_HM group versus the CP_CM group (figure 2E). Our IPA analysis further showed that genes in the Coordinated Lysosomal Expression and Regulation (CLEAR) pathway involving lysosome-associated processes, such as autophagy, phagocytosis, and immune response were also more accessible/had relaxed chromatin state in the HP_HM group. Similar to our RNA-seq analysis (figure 2D), we found genes involved in DNA Damage pathways showing increased accessibility in HP_HM group compared to the CP_CM group. Such pathways included the DNA damage-induced 14-3-3σ Signaling, Melatonin Degradation I, and Super Pathway of Melatonin Degradation (figure 2E). It is well known that Melatonin Degradation causes oxidative DNA damage [39]. To further support our RNA-seq results further, we found genes impacting mitochondrial functionality were found have reduced accessibility in the HP_HM group versus the CP_CM group similar to the low expression of such genes in the RNA-seq analysis.

Above studies suggest that the upregulation observed in pathways relating to DNA damage, inflammation, tissue damage and downregulation in homeostatic-stress signaling and wound healing pathways observed in the intestinal tissues of hydromorphone exposed group (HP_HM) versus control group (CP_CM) as seen in the RNA-sequencing analysis, is partially due to the changes in chromatin accessibility of the respective genes involved in the corresponding pathways.

### Cross-fostering hydromorphone pups to control mothers attenuates hydromorphone-associated small intestinal gut microbiome change

To delineate the relative contributions of prenatal versus postnatal factors in offspring gut microbiome development, a cross-fostering paradigm was developed. Hydromorphone-exposed pups were cross-fostered to control dams (HP_CM), and control pups were cross-fostered to hydromorphone-treated dams (CP_HM) (Supplementary Figure 3A). This approach enables differentiation of prenatal influences from postnatal maternal and environmental effects by comparing offspring microbiomes raised by dams from either the same or opposite treatment groups [40]. Pups were cross fostered at postnatal day 3 until offspring were 3 weeks of age. To determine the impact of cross fostering on offspring gut-microbial composition, stool samples from HP_HM, HP_CM, CP_CM, or HP_HM offspring were subjected to 16S rRNA sequencing. Alpha diversity (Shannon index) of stool samples was significantly higher in HP_CM offspring relative to HP_HM offspring (Fig 3A), and cross fostering resulted in significant changes in gut microbial composition per the Bray-Curtis dissimilarity index (PERMANOVA: F = 3.666, R2 = 0.314, *p* = 0.012) between HP_CM and HP_HM offspring (Fig 3B). At the phylum level, HP_CM pups had significantly increased mean relative abundance of *Proteobacteria* (p=0.029) (Fig 3C). At the genus level, cross fostering hydromorphone pups to control mothers resulted in significant differences in 7 genera (Fig 3D-J). Among these, cross fostering rescued decreases in *Romboutsia* and *Lachnospiracea A2* in HP_HM offspring and further increased the relative abundance of *Parasuterella* and *Eisenbergiella* in the HP_CM group relative to HP_HM (Fig 3D-G). Additionally, cross fostering was also able to rescue significant increases in *Lachnospiraceae UCG-006* and *Lachnoclostridium* in HP_HM relative to HP_CM, as well as decrease the relative abundance of *Enterorhabdus* (Fig 3H-J).

**Figure 3.**
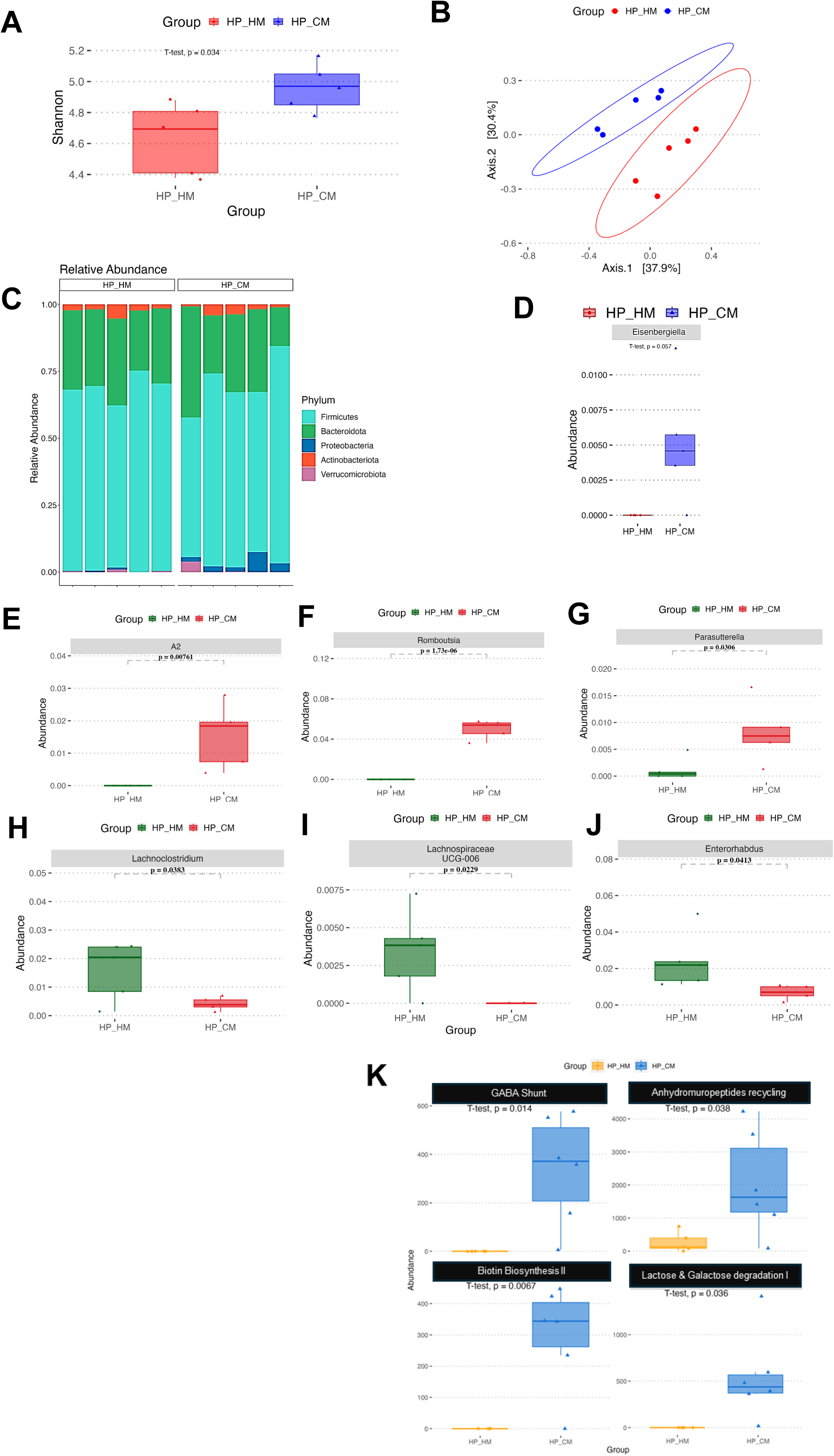
Cross-fostering hydromorphone pups to control mothers attenuates hydromorphone-linked intestinal gut dysbiosis (n=5/group) **(A)** Shannon diversity index in HP_HM (red) versus HP_CM (blue); box = IQR, whiskers = 1.5×IQR, line = median; *p* < 0.05 by two-sided *t*-test. **(B)** Principal coordinates analysis of Bray–Curtis distances; each point is a sample and ellipses denote 95% confidence intervals for each group. **(C)** Stacked barplot showing phylum-level relative abundances in each sample; phyla are colored as indicated in the legend. **(D–J)** Relative abundance of selected genera that differ significantly between HP_HM and HP_CM: **(D)** *Eisenbergiella;* **(E)** *A2;* **(F)** *Romboutsia;* **(G)** *Parasutterella;* **(H)** *Lachnoclostridium;* **(I)** *Lachnospiraceae_UCG-008;* **(J)** *Enterorhabdus.* Boxplots are plotted as in (A); *p*-values by unpaired two-sided *t*-tests and indicated above each panel. **(K)** PICRUST functional analysis. Boxplots showing relative abundance of four representative pathways; *p*-values by Bonferroni-adjusted pairwise *t*-tests.

We further explored our 16S rRNA sequencing data to understand the impact of cross fostering on the functional capabilities of microbial communities in the gut microbiome. The microbial metagenome was inferred using the PICRUST analysis and functional profiles were classified according to KEGG pathways to better understand specific alterations in microbial pathways. We observed changes in bacterial taxa with significant differences in the metabolic profile of many distinct pathways (Supplementary Figure **3B**). Interestingly, pathways relevant for Biotin Biosynthesis and for Short-chain Fatty acid synthesis (SFCAs) such as the Gamma-Aminobutyric Acid Shunt (GABA Shunt), Lactose and Galactose Degradation Pathway, Anhydromuropeptides Recycling were earlier found to be significantly reduced following prenatal hydromorphone exposure (Fig 1K). Cross fostering hydromorphone pups to control mothers rescued the decrease in six of these pathways namely, Biotin biosynthesis II, Glycerol degradation to butanol, 4-aminobutanoate (GABA) degradation V, Anhydromuropeptides recycling, Lactose and Galactose degradation I, and Methanogenesis from Acetate in the HP_CM group relative to HP_HM (Fig 3K). Additionally, the analysis revealed an increase in signaling pathways such as thiazole biosynthesis, tRNA processing, super-pathways of glycolysis and Entner-Doudoroff, hexitol degradation (bacteria), polyamine biosynthesis I/II, thiamin diphosphate biosynthesis, and TCA cycle VII (acetate-producers) were predicted to be upregulated in HP_CM relative to HP_HM offspring. We then performed Bugbase analysis that revealed that cross fostering hydromorphone pups to control mothers was able to rescue significant increase in Facultatively Anaerobic bacteria in HP_HM relative to HP_CM group (Supplementary Table 1B).

Cross fostering control pups to hydromorphone mothers had limited effects on gut microbiome composition. No differences in alpha diversity (p=0.56), beta diversity (PERMANOVA: F = 1.291, R2 = 0.114, *p* = 0.273), or differentially enriched phyla were noted between CP_CM and CP_HM groups (Supplementary Fig 4A-C). At the genus level, CP_HM offspring exhibited an increased relative abundance of *Enterorhabdus* and *Turicibacter* relative to CP_CM (Supplementary Fig 4D-E). Changes in these bacterial taxa were associated with significant differences in the metabolic profile of 4 distinct pathways per PICRUST analysis (Supplemental Fig 4F). Pathways related to the adenine/adenosine salvage, purine ribonucleosides degradation, super pathway of pyrimidine deoxyribonucleoside degradation, and tRNA processing were predicted to be upregulated in CP_HM offspring relative to CP_CM. In summary, cross fostering control pups to hydromorphone mothers appears to have limited effects on the gut microbiome.

Collectively, these findings demonstrate that cross fostering prenatally exposed hydromorphone pups to control mothers partially restores the gut microbial composition. Specifically, the relative abundance of beneficial taxa such as *Romboutsia* and *Lachnospiraceae A2* was rescued, coinciding with upregulation of the GABA degradation pathway and other key biosynthetic pathways. In contrast, the cross-fostering intervention prevented the overrepresentation of potentially pathogenic genera, including *Lachnoclostridium* and *Enterorhabdus*.

### Cross-fostering hydromorphone pups to control mothers attenuates hydromorphone-associated small intestinal damage

Our laboratory has previously demonstrated that opioid exposure causes epithelial injury, apical villous expulsion and inflammatory cell influx in intestinal lumen in both male and female mice [29]. We hypothesized that prenatal hydromorphone exposure damages offspring small intestinal lumen which is attenuated when hydromorphone pups are cross-fostered to control mothers. Distal intestinal tissues were collected from pups belonging to CP_CM, HP_HM, and HP_CM groups. The ileal tissue sections were stained with hematoxylin and eosin (H&E) and scored using the established criteria (Figure 4B). Contrary to the control samples (CP_CM), prenatal hydromorphone exposure associated with epithelial injury to the intestinal villus in pups belonging to the HP_HM group (Fig 4A). Dramatically, prenatal hydromorphone exposure showed apical villus expulsion into lumen, causing substantial tissue damage when compared to CP_CM group. Interestingly, cross fostering prenatally hydromorphone exposed pups with control mothers (HP_CM group) attenuated the damage observed in the intestinal villus and maintained the intestinal lumen architecture (Fig 4A-C). A Pearson’s correlation analysis was performed between gut bacteria identified in Figure 1 (16 sRNA-seq) and intestinal damage score analysis (Figure 4C), to determine whether microbial dysbiosis observed following prenatal hydromorphone exposure (HP_HM) was related to the intestinal tissue damage. There was negative correlation between the damage scores and the gut bacteria-*Bifidobacterium, A2,* and *Romboutsia*, while a positive correlation was observed between damage score and the gut bacteria namely-*[Eubacterium] xylanophilium group*, *Lachnospiraceae UCG-006*, and *Lachnoclostridium* (Figure 4D). We then performed the Pearson’s correlation analysis to explore if the less damage observed while cross fostering prenatally hydromorphone exposed pups with control mothers (HP_CM group) was related to changes in gut microbiome observed in Figure 3. We observed negative correlation between the damage scores and the gut bacteria-*Romboutsia, A2, Parasuterella,* and *Eisenbergiella*, while a positive correlation was observed between damage scores and the gut bacteria namely-*Lachnoclostridium, Lachnospiraceae UCG-006*, and *Enterorhabdus*. (Figure 4E).

**Figure 4.**
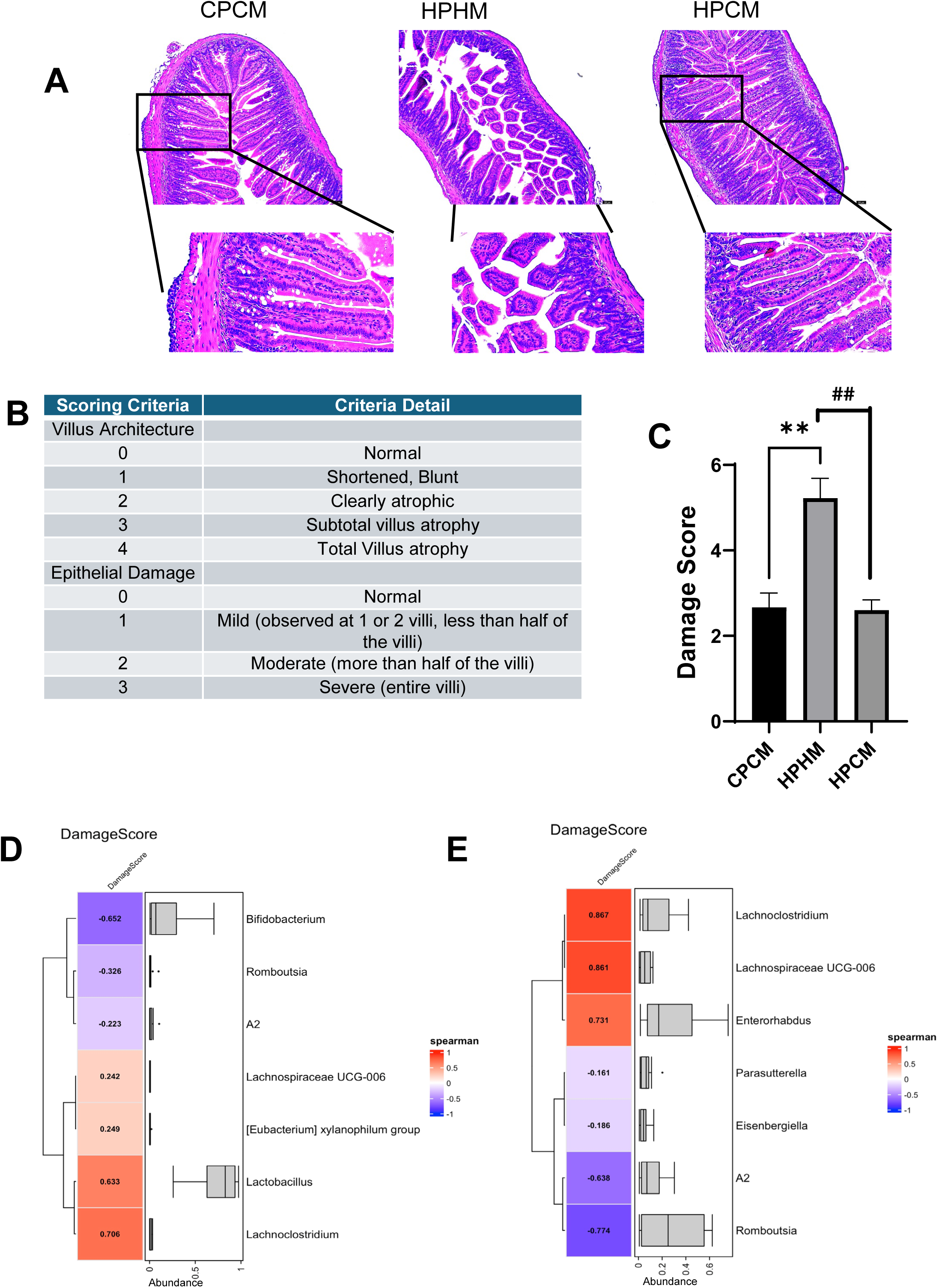
Cross-fostering hydromorphone pups to control mothers attenuates hydromorphone-associated small intestinal damage (n = 6 each). **(A)** Representative H&E-stained small intestinal sections from CP_CM, HP_HM, and HP_CM. Scale bar: 50 µM. **(B)** Histopathological damage scoring guideline for H&E-stained sections. **(C)** Graphs showing histopathological damage score in intestinal sections in different treatment groups. Data represented as bar plots with Standard error of mean (SEM). Data was analyzed by one-way ANOVA with Student T-Test (p<0.001) **(D)** Spearman’s Correlation analysis between Intestinal Damage Score and bacterial taxa between HP_HM and CP_CM. Red indicated positive correlation and Blue indicated negative correlation. **(E)** Spearman’s Correlation analysis between Intestinal Damage Score and bacterial taxa between HP_HM and HP_CM. Red indicated positive correlation and Blue indicated negative correlation.

These data suggest that cross fostering prenatally exposed hydromorphone pups to control mothers is able to attenuate the hydromorphone-associated intestinal damage observed in HP_HM group, possibly due to the rescuing of the gut bacteria, which is inclusive of *Romboutsia and A2,*

### Cross-fostering hydromorphone pups to control mothers attenuates hydromorphone-associated transcription and chromatin profiles corresponding to intestinal tissue damage and reduced homeostatic-stress mechanisms

We hypothesized that the attenuation in damage to the distal intestinal tissue observed while cross-fostering hydromorphone pups to control mothers (HP_CM group) is due to changes in gene transcription impacting pathways involved in tissue repair, inflammation and cellular damage versus HP_HM group. To determine the transcriptional underpinnings corresponding to the impact of cross fostering on offspring gut, bulk RNA-seq analysis was performed on small intestinal tissue of 3-week-old HP_CM and HP_HM groups. Applying the adjusted p-value <0.05 & |log2FC| >0.5 was too stringent and so we resorted to using p-value <0.05 & |log2FC| >0.5. Differential gene expression analysis revealed 139 genes were significantly upregulated and 112 genes were downregulated in HP_CM group relative to HP_HM (Fig. 5A). To understand the changes observed in HP_CM group versus HP_HM group, we applied the IPA analysis platform to identify the biological pathways and potential mechanisms involved. The macrophage alternative activation signaling pathway, which helps in wound healing and repair, was the top pathway rescued in the HP_CM group relative to HP_HM group, per IPA analysis on unadjusted *p*<0.05, |LFC|>1.0 gene set (Fig 5C). Additionally, homeostatic-stress-stress response pathways involved in gut homeostasis such the Sirtuin Signaling, IL-7 signaling pathway, Glucocorticoid Receptor Signaling, B cell Development, B cell receptor Signaling, Phagosome formation pathway genes were upregulated versus HP_HM group. We also found genes impacting Granzyme A Signaling and Mitochondrial dysfunction were upregulated in HP_CM group versus HP_HM. Interestingly, we found downregulation of several pathways in HP_CM group that were implicated in Oxidative Phosphorylation, Ferroptosis, Th2 Signaling, Neutrophile Extracellular Trap Signaling, and Systemic Lupus Erythematous in B cell Signaling, as compared to HP_HM group. Our RNA-seq analysis therefore revealed that cross fostering prenatal hydromorphone pups to control mothers (HP_CM group) showed pathways enriched in repair and stress-adaptive response and downregulation of inflammatory and tissue damage pathways versus HP_HM.

**Figure 5.**
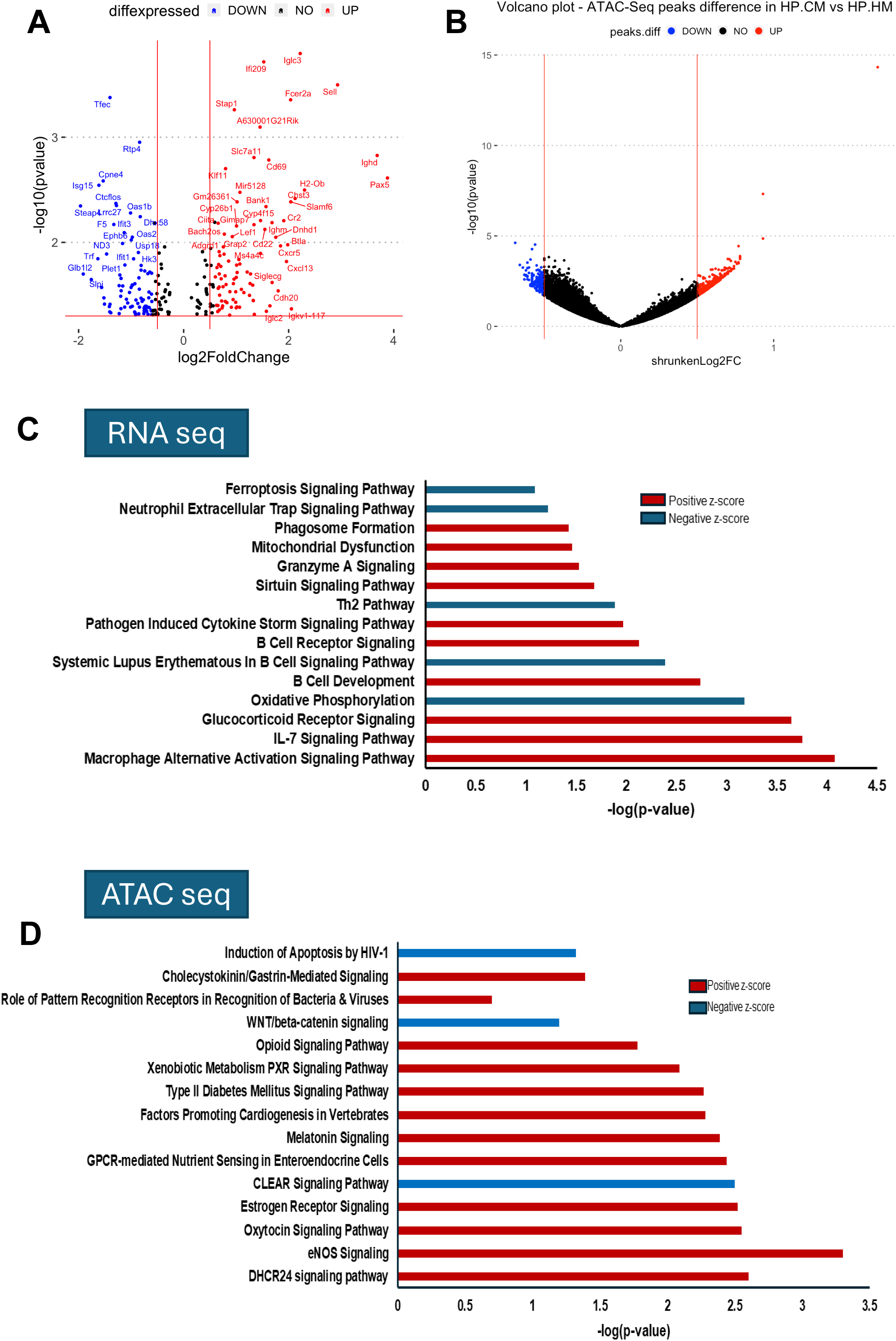
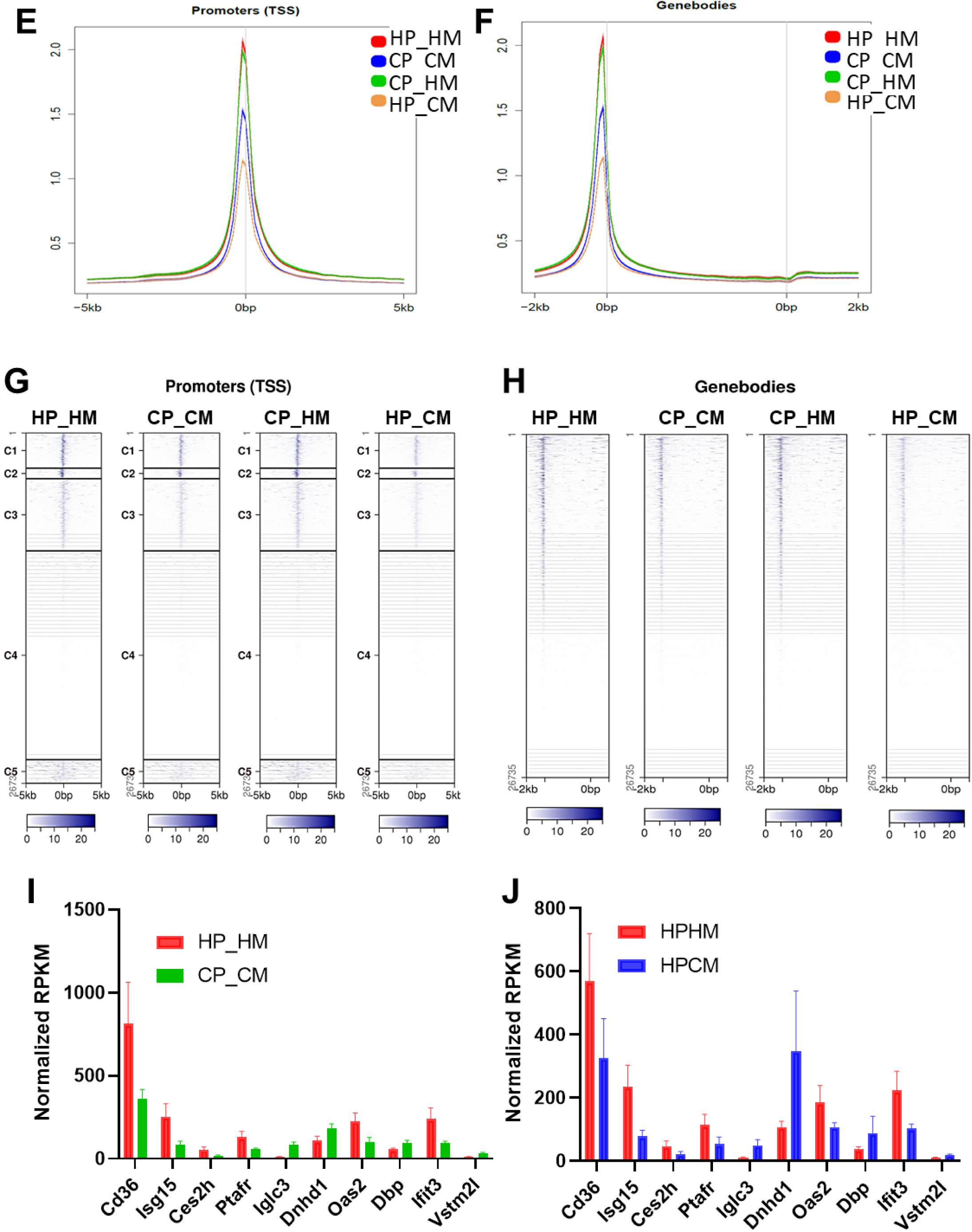
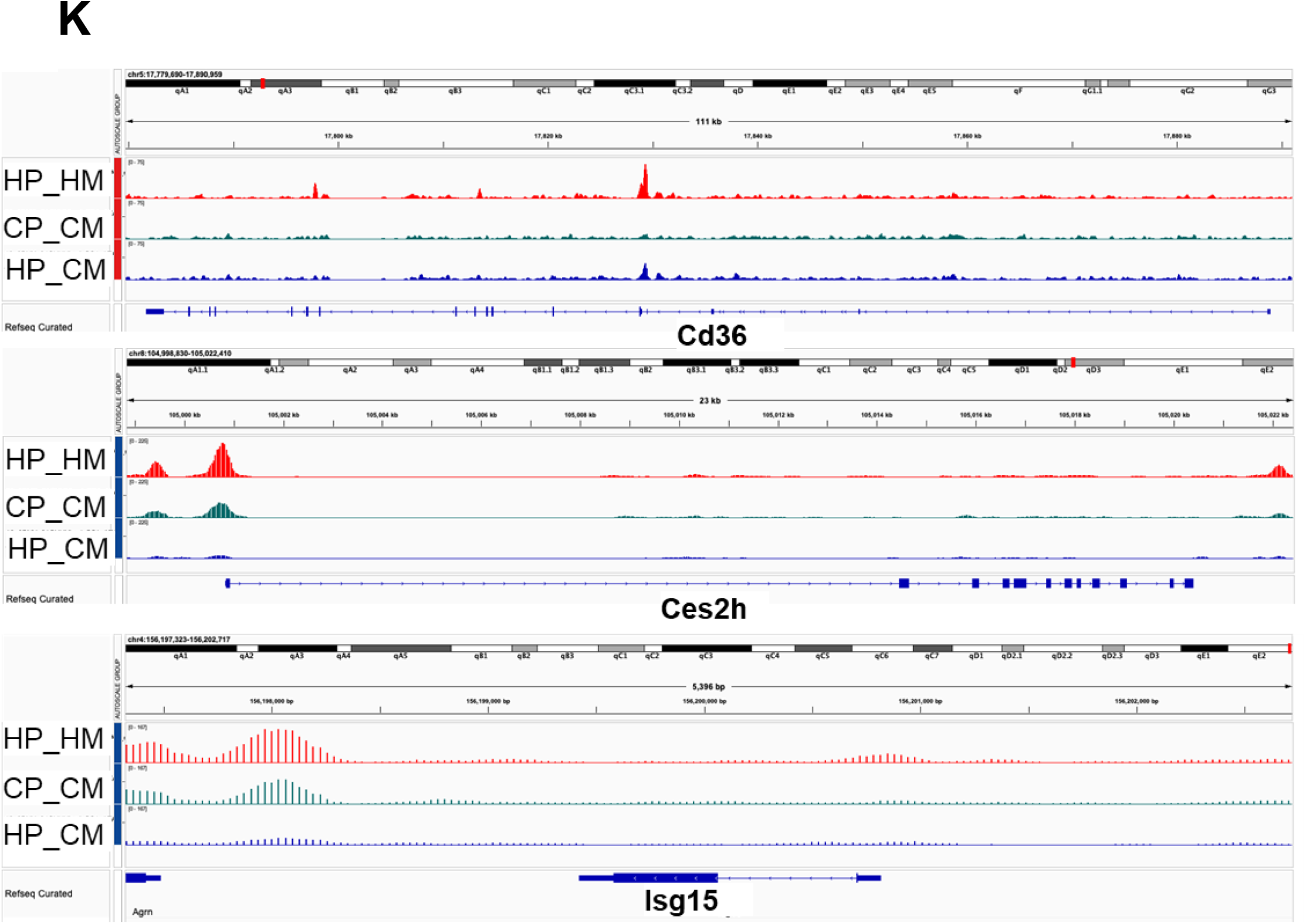
Cross-fostering hydromorphone pups to control mothers attenuates hydromorphone-associated transcription and chromatin profiles corresponding to intestinal tissue damage (n=5/group). **(A)** Volcano plot of RNA-seq differential gene expression (HP_CM vs HP_HM): log₂ fold change vs –log₁₀(*p*). Genes with *p*<0.05 and |LFC|>0.5 are colored red (up in HP_CM) or blue (down); selected DEGs are labeled. **(B)** Volcano plot of ATAC-Seq differential peak accessibility (HP_HM vs HP_CM). Each point represents a consensus peak, colored by significance (*p*<0.05, |shrunkLog₂FC|>0.5) as in (A). Thresholds are shown by dashed lines. **(C)** Top enriched canonical pathways from IPA based (from RNA-seq between HP_CM vs HP_HM) on the unadjusted *p*<0.05, |LFC|>1.0 gene set. Bars denote –log₁₀(*p*); red, positive *z*-score (predicted activation in HP_HM); blue, negative *z*-score (predicted inhibition). Because FDR filtering returned too few genes for reliable pathway calls, unadjusted *p* was used to expand the gene set. **(D)** Top enriched canonical pathways from IPA based on the unadjusted *p*<0.05, of genes with differential ATAC peaks in HP_CM versus HP_HM. Bars denote –log₁₀(*p*); red, positive *z*-score (predicted activation in HP_CM); blue, negative *z*-score (predicted inhibition). Because FDR filtering returned too few genes for reliable pathway calls, unadjusted *p* was used to expand the gene set. **(E)** Enrichment of ATAC-seq signal around the transcription start site (TSS) to gauge gene promoters. Data is merged from n=5 samples. Aggregated enrichment of genes that are induced by hydromorphone treatment. **(F)** Enrichment of ATAC-seq signal in gene bodies. Data is merged from n=5 samples. Aggregated enrichment of genes that are induced by hydromorphone treatment. **(G)** Representative heatmap profile of differentially accessible peaks for promoters between HP_HM, CP_CM, CP_HM, and HP_CM. **(H)** Representative heatmap profile of differentially accessible peaks for gene bodies between HP_HM, CP_CM, CP_HM, and HP_CM. **(I)** Normalized RPKM gene expression for candidate genes in HP_HM versus CP_CM. **(J)** Normalized RPKM gene expression for candidate genes in HP_CM versus HP_HM. **(K)** Representative traces of ATAC-seq in various comparison groups CP_CM, HP_HM, HP_CM for candidate genes Cd36, Ces2h, Isg15.

We reasoned that the alterations in chromatin accessibility of associated genes in the ileal tissues of cross-fostered hydromorphone pups to control mothers (HP_CM group) is responsible for the changes in gene expression impacting cellular inflammatory, homeostatic-stress, and cellular damage signaling pathways when compared to the HP_HM group. To understand the chromatin architecture status of these samples, we resorted to ATAC-sequencing analysis (Fig 5B). We studied the ATAC-seq peaks at beta-Actin gene loci for control between the comparison groups HP_CM and HP_HM (Supplementary Fig 2). As expected, we observed that the chromatin accessibility at the beta-Actin gene loci did not change between the HP_CM versus HP_HM comparison groups. We further utilized the Ingenuity Pathway Analysis platform on the accessible chromatin regions identified through ATAC-seq for the HP_CM and HP_HM groups to identify biological pathways and potential mechanism involved. Our analysis showed that chromatin accessibility to several genes involved in homeostatic-stress or gut health aiding pathways showed “open chromatin”, such as Melatonin Signaling, DHCR24 Signaling pathway, eNOS Signaling, Oxytocin Signaling, Estrogen Receptor Signaling, GPCR-mediated Nutrient Sensing in Enteroendocrine Cells, Factors Promoting Cardiogenesis in Vertebrates, Xenobiotic Metabolism PXR Signaling, Cholecystokinin/Gastrin-Mediated Signaling (Fig 5D). Additionally, we found increased accessibility to various genes implicated in pathways involved in cell death and inflammation such as Induction of Apoptosis by HIV-1 signaling, Coordinated Lysosomal Expression and Regulation, or CLEAR Signaling, and WNT/beta-Catenin Signaling, showed a more “closed chromatin” conformation down regulated in HP_CM group in comparison to HP_HM group (Fig 5D).

To test if prenatal hydromorphone exposure (HP_HM) affects the chromatin status and whether cross fostering hydromorphone pups to control mothers (HP_CM) rescues these changes, we studied the changes in the chromatin accessibility both around the Transcription Start Site (TSS) of genes as well as at the Gene Bodies (Fig 5E-H). Gene Promoters around the TSS span the regulatory regions and chromatin accessibility around them is an important indicator of gene expression status. It is worth noting that gene regulatory regions can span many kilobases either upstream or downstream of the TSS into Gene Bodies [41]. We observed a striking diffused relaxation of chromatin following prenatal hydromorphone exposure (HP_HM group) (Fig 5E-H). Interestingly, we found that cross-fostering hydromorphone pups to control mothers (HP_CM) rescued the chromatin accessibility alterations observed both at the Gene Promoters around the TSS and Gene Bodies within the HP_HM group. Note that the chromatin relaxation observed in the HP_HM group was more similar to the chromatin relaxation status observed when control pups were cross-fostered to hydromorphone mothers (CP_HM group). On the other hand, we found that cross-fostering hydromorphone pups to control mothers (HP_CM group) attenuates the chromatin relaxation observed in HP_HM group and makes the chromatin more “closed/tight” both at the Gene Promoters around the TSS and Gene Bodies (Fig 5E-H), if not resembling the CP_CM group.

We previously observed upregulation in pathways relating to inflammation, tissue damage and downregulation in homeostatic-stress signaling pathways in the intestinal tissues of hydromorphone exposed group (HP_HM) versus control group (CP_CM) (Fig 2D-E). We took candidate genes from our RNA-seq analysis implicated in these pathways and observed the gene expression analysis in comparison groups HP_HM versus CP_CM and HP_CM versus HP_HM (Fig 5I-J). Genes implicated in gut disruption and inflammation such as *Cd36, Isg15, Ces2h, Ptafr, Oas2,* and *Ifit3* were found to be upregulated in HP_HM group versus CP_CM group, while the same genes were found to be downregulated in HP_CM group versus HP_HM group comparison. Interestingly, genes implicated in gut barrier function and integrity such as D*bp, Vstm2l,* and *Dnhd1* were found to be downregulated in HP_HM groups versus CP_CM group, while the same genes were found to be upregulated in HP_CM group when compared to HP_HM group (Fig 5I-J). We then studied the chromatin accessibility status of these genes between the comparison groups of HP_HM, CP_CM, and HP_CM. We found that the ATAC-seq peaks in genes such as the *Cd36, Isg15,* and *Ces2h*, was more in HP_HM group versus CP_CM group (Fig 5K). The ATAC-seq peaks in the HP_CM group for these genes were also less than the HP_HM group. This indicated that the chromatin architecture is more “tighter/closed” at the indicated gene loci which correlated to less expression in CP_CM and HP_CM group when compared to HP_HM group (Fig 5I-J).

Collectively, these data provide evidence that prenatal hydromorphone exposure is associated with gut dysbiosis and intestinal tissue damage, and cross-fostering hydromorphone pups to control mothers partly attenuates hydromorphone-associated gene expression and chromatin profiles corresponding to intestinal tissue damage, oxidative stress, inflammation, and reduced homeostatic-stress mechanisms, which might explain the reduced intestinal damage observed in HP_CM group.

### Fecal microbiota transplantation with feces from hydromorphone exposed dams show increased intestinal tissue damage

To further validate if the alterations in gut microbiome can cause damage to the intestinal tissue, we gavaged 3-week-old naïve offspring, with either fecal samples from hydromorphone exposed dams, or, those from the saline treated dams in the same ratio with the hydromorphone exposed fecal sample (Fig 6A). As expected, FMT from hydromorphone exposed dams resulted in significant damage to the distal ileal tissues of naïve 3-week-old offspring (Fig 6B-C). Interestingly, FMT from saline and hydromorphone exposed dams (1:1 ratio) rescued the damage associated with hydromorphone exposed dams (Fig 6B-C). This indicated that rescuing gut morphology owing to the damage associated with opioid exposure is possible provided the presence of potentially relevant microbiota or the reduced presence of harmful bacteria. Our previous analysis indicated that cross-fostering hydromorphone pups to control mothers (HP_CM) showed increased presence of *Romboutsia* and *A2,* which negatively correlated to the intestinal tissue damage and reduced presence of *Lachnoclostridium, Enterorhabdus,* and *Lachnospiraceae UCG-006,* which positively correlated to the intestinal tissue damage (Fig 4D-E). Further, to identify significant associations between gut microbiota and the ileal transcriptome, we performed a correlation analysis where presence of the gut microbe-*Romboutsia* and *Lachnospiraceae A2* (known to produce SCFAs and other favorable metabolites required for gut health) which were recovered in the HP_CM group, were correlated to DGEs in our RNA-seq analysis comparing HP_CM versus HP_HM (Fig 6D). We found that genes such as Dbp, Dnhd1 (gut development [42]), Vstm2l (protects cells from ferroptosis (a form of programmed cell death), helping maintain mitochondrial function and cell viability [43]), Cd69 (suppresses excessive inflammation and offers gut barrier protection [44]), Acot3 (support metabolism and prevent lipotoxicity [45]), etc. were positively correlated with the presence of either *Romboutsia* or *A2* (Fig 6E).

**Figure 6.**
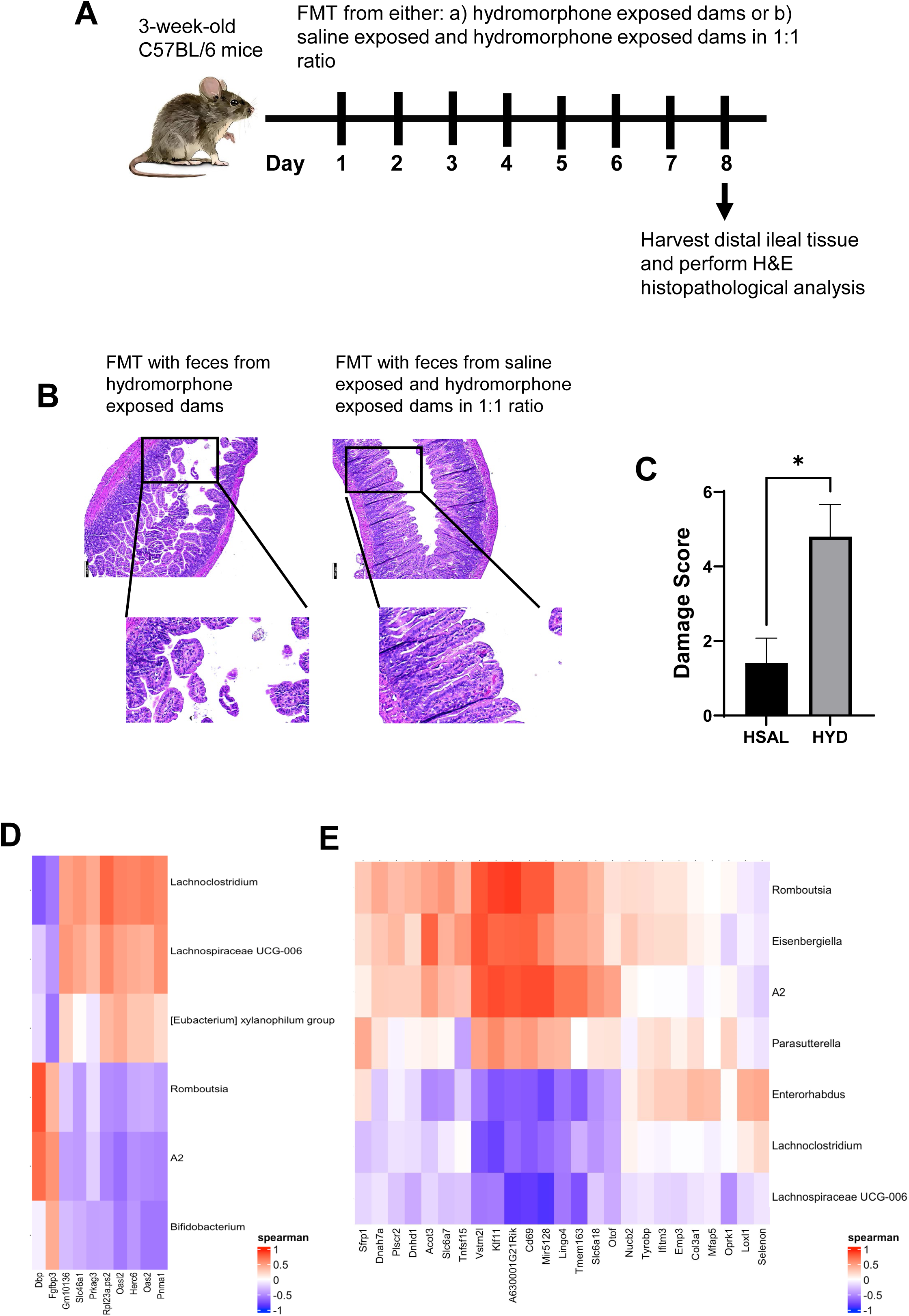
Fecal microbiota transplantation with feces from hydromorphone exposed dams show increased intestinal tissue damage. (A) Schematic for FMT with stool samples from opioid dams or opioid and saline exposed dams into 3-week-old naïve C57BL/6 mice. (B) Representative H&E-stained small intestinal sections from: left panel) pups with FMT from hydromorphone exposed dams, right panel) pups with FMT from saline exposed and hydromorphone exposed dams in 1:1 ratio. Scale bar: 50 µM. (C) Graphs showing histopathological damage score in intestinal sections in different treatment groups. Data represented as bar plots with Standard error of mean (SEM). Data was analyzed by one-way ANOVA with Student T-Test (p<0.05) (D) Spearman’s Correlation analysis between bacterial taxa between HP_HM versus CP_CM with DEG |log2FC| ≥0.05 & p-value ≤0.05. Upregulated genes are indicated in Red while Blue indicated genes that are downregulated. (E) Spearman’s Correlation analysis between bacterial taxa between HP_CM versus HP_HM with DEG |log2FC| ≥0.05 & p-value ≤0.05. Upregulated genes are indicated in Red while Blue indicated genes that are downregulated.

Together, these data suggest that alterations in gut microbiome (brought about by FMT from hydromorphone exposed dams) can lead to increased damage to the intestinal tissue.

## Discussion

Prenatal opioid exposure, identifying altered methylation patterns in genes involved in neurodevelopment, synaptic function, and drug addiction [46–48]. To our knowledge, this is the first study that utilizes ATAC-sequencing analysis in ileal tissues to illustrate the impact of opioid exposure on chromatin accessibility on a global scale in offspring. Previous chromatin accessibility studies have been limited to the brain in opioid use disorder [51, 52]. We show that prenatal opioid exposure is linked with ileal tissue damage which is mediated by the disruption in gut microbiome in offspring and cross-fostering hydromorphone exposed offspring to control dams rescues the gut disruption, indicating availability of a window of therapeutic intervention. Additionally, our ATAC-seq and RNA-seq analysis suggest that prenatal opioid exposure can potentially alter chromatin accessibility changes resulting in increased expression of genes implicated in inflammation, oxidative stress and reduced homeostatic-stress signaling pathways while cross fostering reduced further increase in inflammatory/damaging pathways and restored wound-healing and repair pathways. Further mechanistic studies are needed to define how opioid exposure drives these chromatin-associated changes.

We developed a clinically relevant model using the iPRECIO 310R infusible mini pump, which provides highly accurate (±5%) and programmable delivery of hydromorphone or saline, closely mimicking clinical opioid exposure patterns [49]. We analyzed microbiome beta-diversity and composition in our control group (CP_CM) and prenatal hydromorphone exposed group (HP_HM) Prenatal hydromorphone reduced α-diversity and shifted β-diversity, with distinct clustering of CP_CM and HP_HM samples. Studies from our lab and others have previously shown that brief [50] and chronic opioid exposure causes gut dysbiosis in offspring [21, 26, 51]. While some studies have reported varying alpha diversity per Chao index [26, 50–52], all consistently reported marked alterations in microbial composition that were evident at weaning [26, 50], during adolescence [52], and into adulthood [51, 52]. Together, this indicates that opioid exposure during critical periods of pregnancy does not merely affect the mother but has profound and lasting consequences for the offspring by interfering with the normal establishment of the gut microbiome during a critical window of development.

Importantly, in our studies, we discovered prenatal hydromorphone exposure resulted in significant differences in 6 genera. The relative abundance of *Bifidobacterium, Romboutsia,* and *A2* was significantly reduced in HP_HM compared to CP_CM, consistent with previous reports [21, 26], showing similar reductions in SCFA-producing bacteria critical for gut homeostasis [53, 54]. We further found that prenatal opioid exposure was linked to increased relative abundance of genera *Lachnoclostridium, [Eubacterium] xylanophilium group*, and *Lachnospiraceae UCG-006* relative to CP_CM group. Published reports have also showed enrichment of these bacteria in prenatally opioid exposed pups [21, 26]. Our PICRUST analysis also supported these results as pathways relevant to metabolite production such as SFCA and Gamma Amino Butyric Acid (GABA) Shunt, Lactose and Galactose Degradation I, and Biotin Biosynthesis II pathways were reduced in prenatally opioid exposed pups versus CP_CM group. Bugbase analysis revealed that prenatal opioid exposure in associated with changes in bacterial composition corresponding to an increase in the predicted relative abundance of facultative anaerobic, am positive, and mobile-element-containing bacteria, and a decrease in the predictive relative abundance of gram negative and biofilm-forming bacteria compared to the CP_CM group. Recent work has demonstrated both in wild-and laboratory-conditions that establishment of an adult-like gut microbiota in house mouse takes around 4 weeks of age [55]. 3-week-old offsprings (where all our experiments were performed), represents the age group where the normal establishment of the gut microbiome takes place. High levels of facultative anaerobic bacteria in the gut lumen are associated with delayed gut development, long-term growth restriction [56] along with dysbiosis and inflammation [57].

Our cross-fostering experiments highlight the roles of biological and environmental factors in shaping the neonatal microbiome. Since cross-fostering occurred at postnatal day 3, early microbial input from biological mothers on days 1 and 2 cannot be fully excluded. Future studies performing cross-fostering immediately after cesarean delivery could better control initial maternal microbial influences. Overall, cross-fostering post-birth offers a therapeutic window to induce lasting changes in the neonatal gut microbiome. Published studies show that the nursing mother and not the birth mother determines the long-term microbial composition of the offspring, and that cross-fostering within the first 48 hours can permanently alter the microbiota and impact disease outcome [58, 59]. Our study shows that pups prenatally exposed to opioids when cross fostered after birth on day 3 to control lactating dams show similar findings. This strategy allows early postnatal microbial intervention, thereby offering a critical window to influence gut establishment and associated health outcomes. Our studies suggest that cross-fostering hydromorphone pups to control mothers (HP_CM group) showed increased α-diversity and significant changes in gut microbial composition versus HP_HM group. Bugbase analysis indicated that HP_CM pups showed rescuing of significant increment in Facultatively Anaerobic bacteria in HP_HM relative to HP_CM group. Furthermore, cross fostering rescue decreases in *Romboutsia* and *A2* in HP_HM pups and also rescue significant increases in *Lachnospiraceae UCG-006* and *Lachnoclostridium* in HP_HM relative to HP_CM, as well as decrease the relative abundance of *Enterorhabdus. Lachnoclostridium* is known to be in higher abundance in several disease contexts such as Crohn’s Disease [60], non-alcoholic fatty liver disease [61], colorectal adeno carcinoma [62], and others. On examining whether changes in gut microbiota conferred changes to the intestinal histology in HP_CM versus HP_HM group, we found that cross-fostering caused significant rescuing of the damage to the ileal tissue in HP_HM group. Interestingly, we found negative correlation between the damage observed in intestinal tissue with *Romboutsia, A2, Eisenbergiella, and Parasutterella* and positive correlation with *Lachnoclostridium, Enterorhabdus*, and *Lachnospiraceae UCG-006*. Our findings are supported by past studies suggesting that alteration in the gut microbiota can damage host tissues [28].

We, therefore, proposed that SCFAs released by the gut bacteria such as Romboutsia, and A2 recovered in the HP_CM pups, could potentially prevent the gut dysbiosis and intestinal damage observed due to prenatal opioid exposure.

We analyzed the chromatin accessibility landscape using ATAC-seq alongside bulk RNA-seq to determine how prenatal opioid exposure affects the intestinal epigenome and whether cross-fostering modulates these changes. ATAC-seq identifies genomic regions where chromatin is “open” and accessible to transcription factors and other regulatory proteins [63–65]. Prenatal hydromorphone exposure led to widespread chromatin relaxation at both gene promoters and gene bodies, whereas cross-fostering to control dams partially reversed this effect. Integration of ATAC-seq with RNA-seq showed concordant pathway-level changes: genes involved in DNA damage, inflammation, oxidative stress, and cell death were upregulated and more accessible in hydromorphone-exposed pups, while pathways supporting immune regulation, wound healing (e.g., macrophage alternative activation), and stress adaptation (e.g., Sirtuin signaling) were suppressed. These findings align with previous brain-focused studies showing that opioid exposure alters chromatin accessibility and neuroinflammatory gene networks [66], but extend such evidence to the gut.

RNA-seq revealed that the macrophage alternative activation signaling pathway, critical for immune regulation and mucosal repair, was strongly upregulated in hydromorphone-exposed pups cross-fostered to control dams (HP_CM). Additional protective pathways supporting gut homeostasis—including Sirtuin signaling [67], IL-7 signaling [68], glucocorticoid receptor signaling [69], B cell development and receptor signaling [70], and phagosome formation [71]—were also enriched, while pathways associated with oxidative stress and tissue damage (oxidative phosphorylation, ferroptosis, Th2 signaling, neutrophil extracellular trap formation, and systemic lupus erythematosus in B cell signaling) were downregulated compared with hydromorphone pups nursed by hydromorphone dams (HP_HM). Granzyme A signaling was also increased in the HP_CM group. ATAC-seq supported these transcriptional trends: genes mediating stress-adaptation and barrier protection (e.g., melatonin, DHCR24, eNOS, oxytocin, estrogen receptor, GPCR-mediated nutrient sensing, xenobiotic metabolism PXR, and cholecystokinin/gastrin pathways) showed increased chromatin accessibility, while pro-inflammatory and cell death pathways (e.g., HIV-1–induced apoptosis, CLEAR signaling, WNT/β-catenin) became less accessible. These epigenetic changes align with the transcriptome findings, indicating partial restoration of protective programs with cross-fostering.

To test whether cross-fostering reverses chromatin changes associated with prenatal opioid exposure, we performed ATAC-seq on ileal tissue. Cross-fostering hydromorphone-exposed pups to control dams restored the chromatin landscape, producing a more closed configuration at both gene promoters and gene bodies compared with HP_HM. We next focused on candidate genes identified by RNA-seq that were linked to inflammation, tissue injury, and gut homeostasis, including Cd36, Isg15, Ces2h, Ptafr, Oas2, Ifit3, Dbp, Vstm2l, and Dnhd1. In HP_HM pups, inflammatory genes (Cd36, Isg15, Ces2h, Ptafr, Oas2, Ifit3) were upregulated, whereas homeostatic genes (Dbp, Vstm2l, Dnhd1) were downregulated. Cross-fostering largely reversed these trends, reducing inflammatory gene expression and increasing gut-protective transcripts. Chromatin accessibility mirrored these changes: promoters of Cd36, Isg15, and Ces2h were more open in HP_HM but became less accessible in HP_CM and CP_CM groups. These findings indicate that cross-fostering not only reshapes the microbiome but also re-establishes protective transcriptional and epigenetic programs.

Finally, our fecal microbiota transplantation (FMT) experiments confirmed that the gut microbiome shaped by prenatal opioid exposure can drive intestinal injury, whereas combining microbiota from control and opioid-exposed dams mitigated this damage. These findings extend prior adult rodent work [12, 14, 15] and highlight cross-fostering as a novel early-life intervention. Overall, prenatal opioid exposure disrupts microbial balance and injures intestinal tissue, accompanied by widespread chromatin opening and induction of inflammatory and stress-response genes. Cross-fostering reverses many of these molecular changes, tightening chromatin and reducing pro-inflammatory gene expression, which likely contributes to the observed tissue protection. Other mechanisms may also influence these effects and warrant future investigation.

In summary, our study provides new evidence for the interplay between prenatal opioid exposure, the gut microbiome, host transcriptome, and chromatin accessibility. Using a clinically relevant murine model, we demonstrate that prenatal hydromorphone exposure disrupts microbial diversity, damages intestinal tissue, and alters gene expression and chromatin architecture in pathways governing inflammation, oxidative stress, and repair. Importantly, cross-fostering to control dams rescued many of these alterations, underscoring the potential of early-life microbial interventions to mitigate opioid-induced harm. Limitations include the use of a single opioid (hydromorphone), assessment at a single developmental time point (weaning), and focus on ileal tissue. Future work should test other opioids, sex as a biological variable, consequence of opioid withdrawal, extend analyses to later developmental stages, and examine additional tissues to fully define the impact of prenatal opioid exposure. Despite these constraints, our findings advance understanding of how prenatal opioid exposure reprograms gut and epigenetic landscapes and point toward actionable strategies for intervention.

---

**Supplementary Figure 1.**
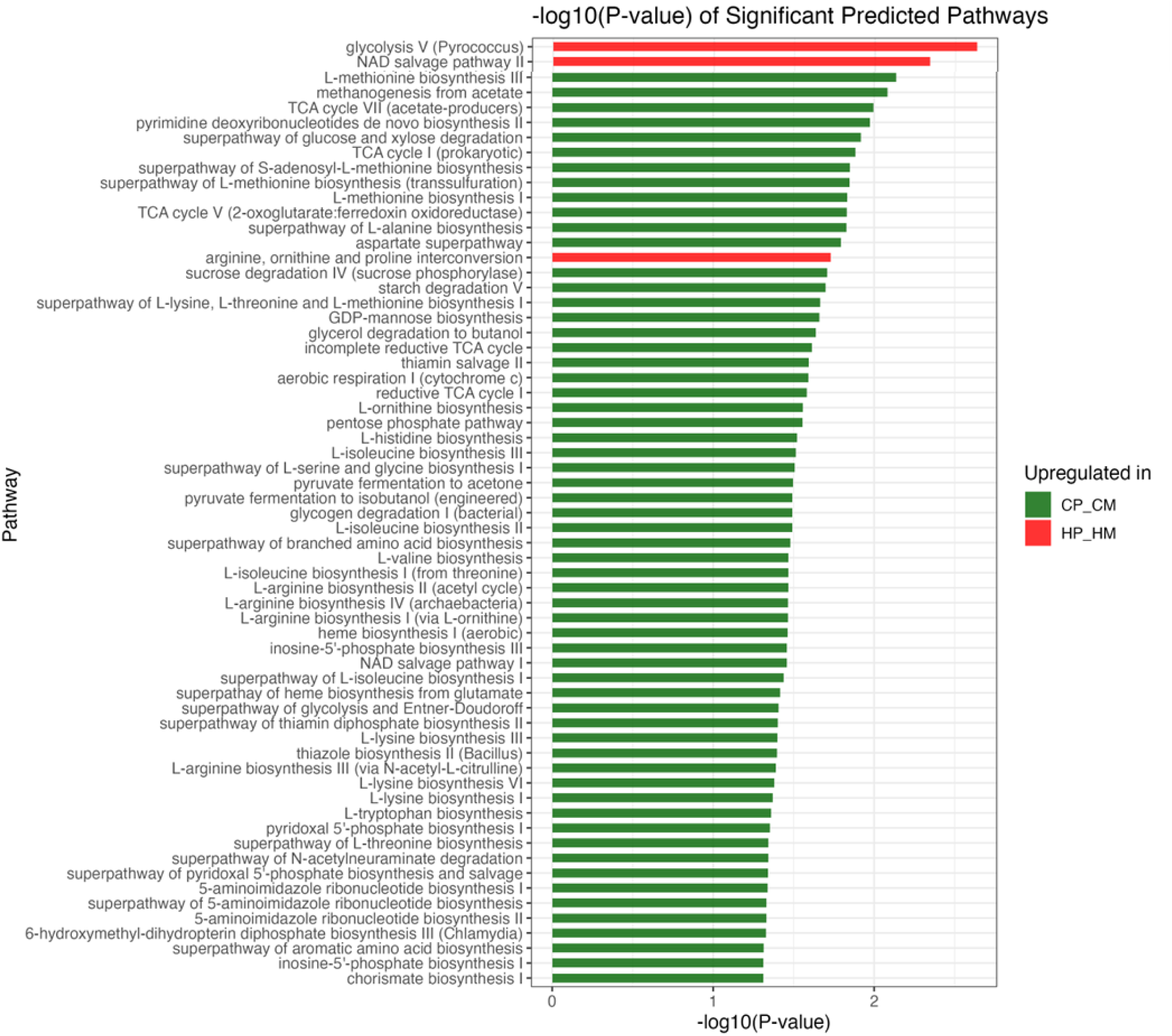
Significant predicted MetaCyc pathways altered in CP_CM vs HP_HM (PICRUSt2 analysis) Horizontal bars show the –log10(*p*) for each pathway that differed significantly between CP_CM (green) and HP_HM (red). Pathways are ordered by decreasing significance (largest to smallest –log10(*p*)). Color indicates the group in which the pathway was predicted to be upregulated. Statistical significance was determined by two-sided Welch’s *t*-tests with Benjamini–Hochberg FDR correction (*p*<0.05).

**Supplementary Figure 2.**
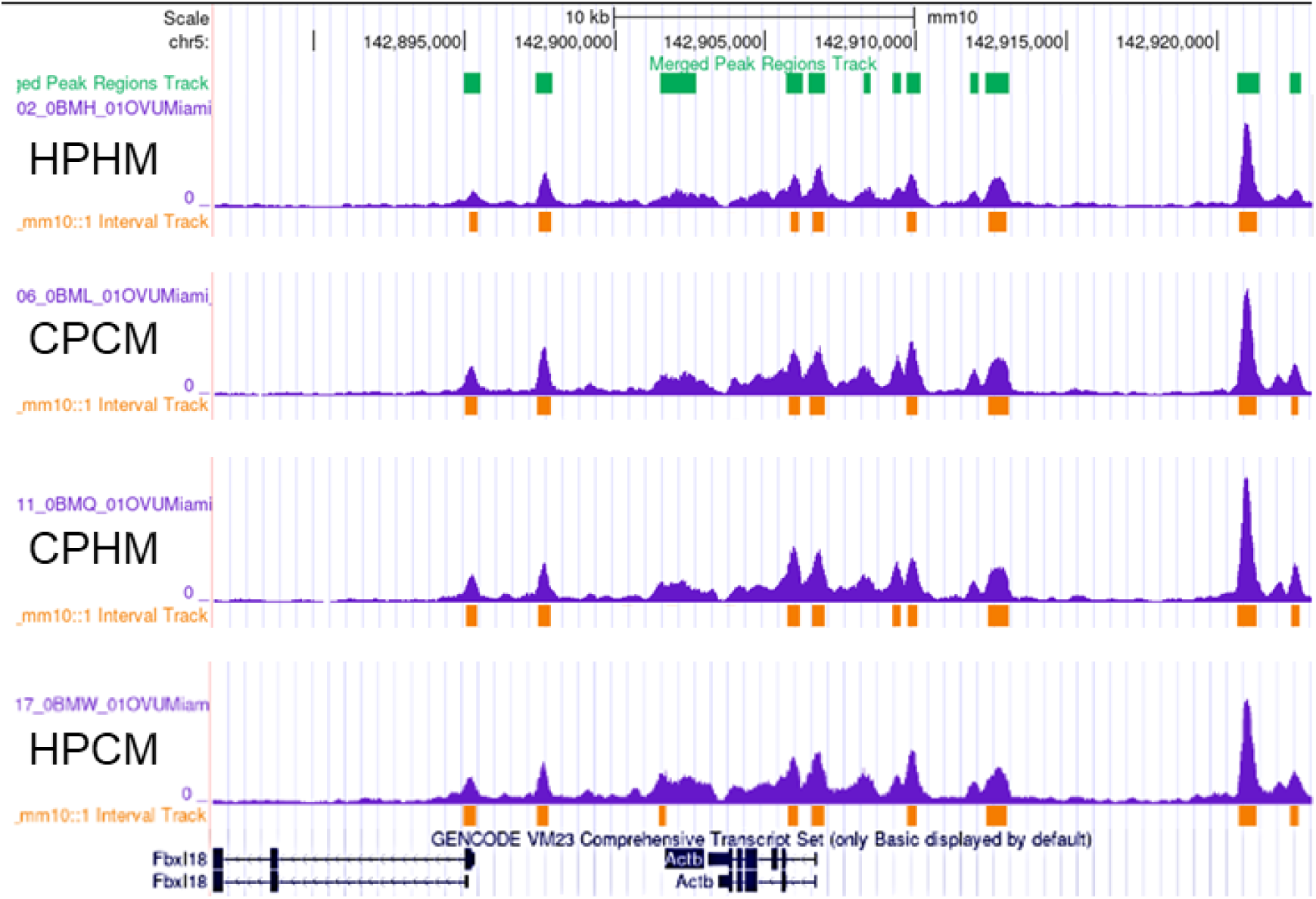
ATAC-seq peaks for housekeeping gene (control loci) beta-actin across all sample groups, HP_HM, HP_CM, CP_HM, CP_CM.

**Supplementary Figure 3.**
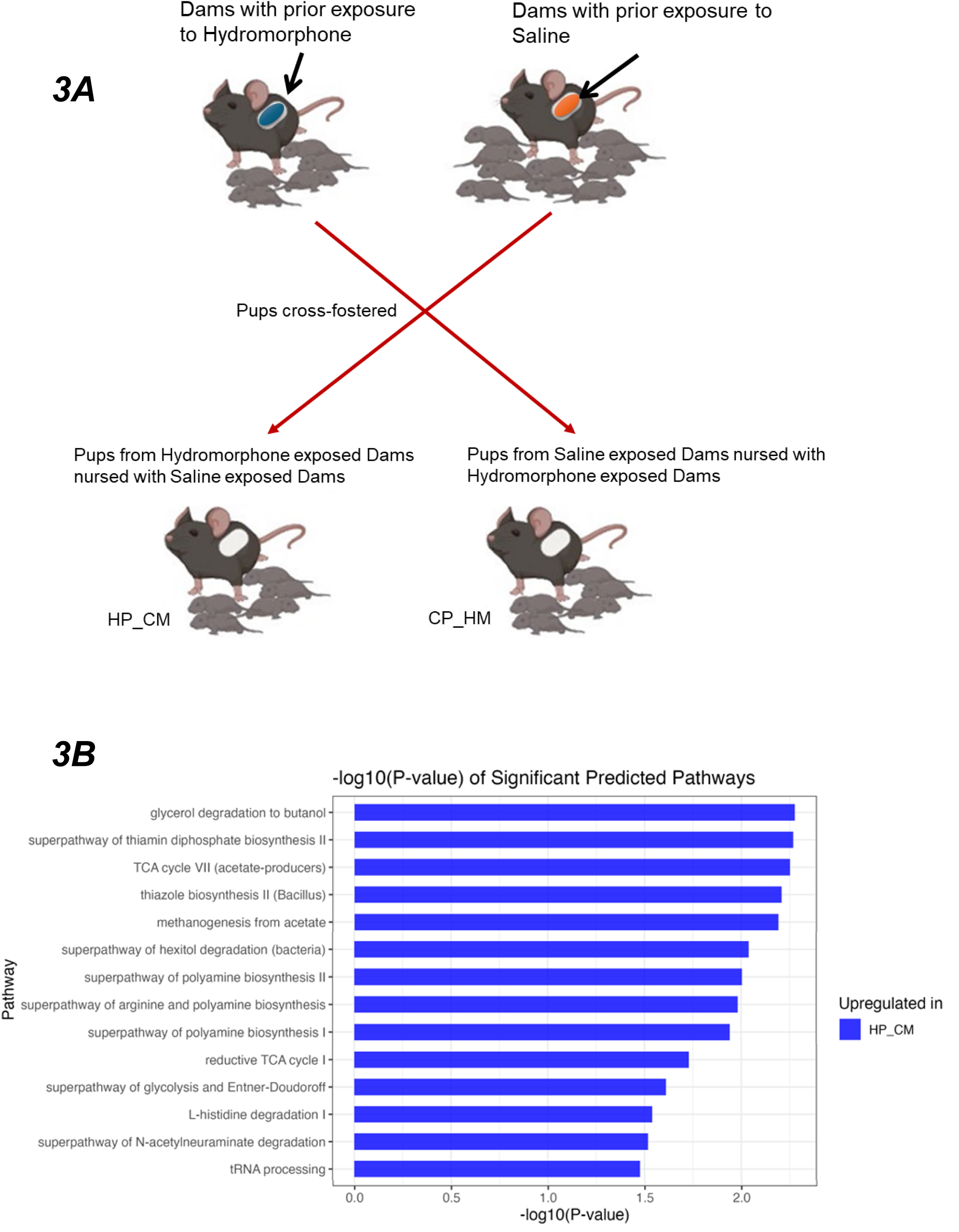
A. *Schematic showing cross fostering strategy.* B. Significant predicted MetaCyc pathways altered in HP_CM vs HP_HM (PICRUSt2 analysis. Horizontal bars show –log₁₀(*p*) for each MetaCyc pathway whose predicted relative abundance differed between HP_CM (blue) and HP_HM (not shown) by unpaired two-sided *t*-test (*p*<0.05). Pathways are ranked from highest to lowest –log₁₀(*p*), indicating the strongest to weakest evidence for differential enrichment in the HP_CM group. Only pathways meeting the significance threshold are displayed; bar length corresponds to the magnitude of –log₁₀(*p*), and color denotes upregulation in HP_CM.)

**Supplementary Figure 4.**
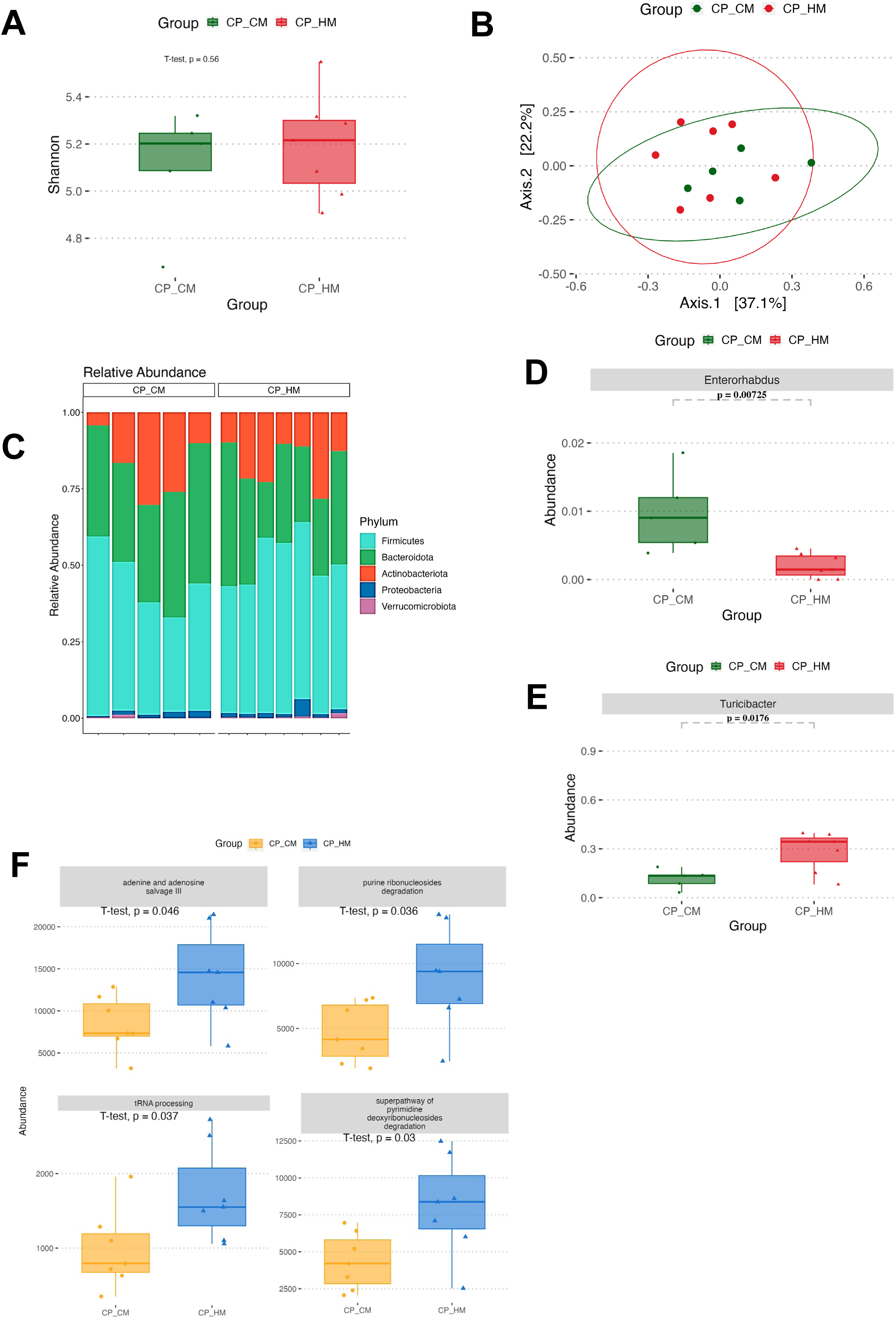
Gut microbiota diversity, composition and key genus differences between CP_CM and CP_HM (n = 5 per group). **(A)** Shannon α-diversity index for CP_CM (green) vs CP_HM (red); box = IQR, whiskers = 1.5×IQR, line = median; *p* by two-sided *t*-test indicated above. **(B)** Principal coordinates analysis (PCoA) on Bray–Curtis distances; each point is a sample and ellipses show 95% confidence intervals for each group. **(C)** Stacked barplots of phylum-level relative abundance in each sample; phyla are color-coded as shown in the legend. **(D)** Boxplot of *Enterorhabdus* relative abundance in CP_CM vs CP_HM; formatting and statistical test as in (A), with *p*-value annotated. **(E)** Boxplot of *Turicibacter* relative abundance in CP_CM vs CP_HM; formatting and statistical test as in (A), with *p*-value annotated. **(F)** PICRUST functional analysis. Boxplots showing relative abundance of four representative pathways; *p*-values by Bonferroni-adjusted pairwise *t*-tests.

## Acknowledgements

National Institutes of Health, **Grant**/Award Numbers: F31DA053795, R01 DA050542, R01 DA047089, R01 DA043252 and R01 DA044582. Hirshberg Foundation for Cancer Research.

## Author contribution statement

SR, SP, and YA conceptualized this project. SP & YA coordinated the experimental design and conducted the project. SP & PS performed data acquisition and analysis of this project. YA provided a draft material on introduction and methods for the manuscript. SP and SR wrote and revised the manuscript. This research was supported by grants from SR. All authors reviewed the results, commented on the manuscript, and approved the final version of the manuscript.

## Data availability statement

The data that support the findings of this study will be openly available in NCBI BioProject ID: PRJNA1353126.

## List of abbreviations

HP_HM: hydromorphone pups nursed by hydromorphone mothers
HP_CM: hydromorphone pups nursed by control mothers
CP_HM: control pups nursed by hydromorphone mothers
CP_CM: control pups nursed by control mothers
ATAC-seq: Assay for Transposase-Accessible Chromatin sequencing

---

**Supplementary Table:**
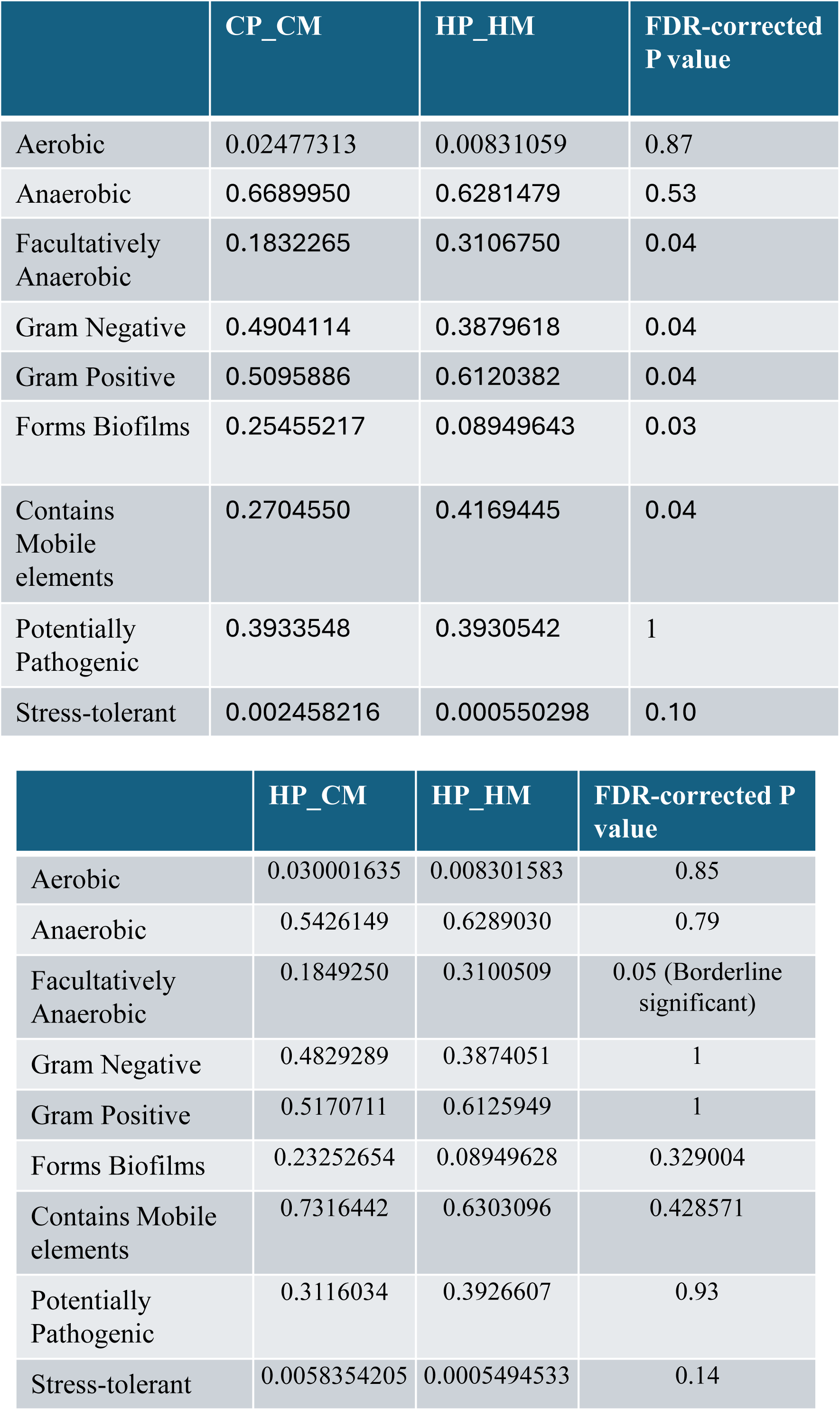
BugBase prediction of the mean relative abundance of predicted high level phenotypes in: Table A) CP_CM and HP_HM fecal samples; Table B) HP_CM and HP_HM fecal samples.

**Supplementary figure 1. Significant pathways altered in CPCM Vs HPHM (PICRUST analysis)**
|  | CP_CM | HP_HM | FDR-corrected P value |
| --- | --- | --- | --- |
| Aerobic | 0.02477313 | 0.00831059 | 0.87 |
| Anaerobic | 0.6689950 | 0.6281479 | 0.53 |
| Facultatively Anaerobic | 0.1832265 | 0.3106750 | 0.04 |
| Gram Negative | 0.4904114 | 0.3879618 | 0.04 |
| Gram Positive | 0.5095886 | 0.6120382 | 0.04 |
| Forms Biofilms | 0.25455217 | 0.08949643 | 0.03 |
| Contains Mobile elements | 0.2704550 | 0.4169445 | 0.04 |
| Potentially Pathogenic | 0.3933548 | 0.3930542 | 1 |
| Stress-tolerant | 0.002458216 | 0.000550298 | 0.10 |

|  | HP_CM | HP_HM | FDR-corrected P value |
| --- | --- | --- | --- |
| Aerobic | 0.030001635 | 0.008301583 | 0.85 |
| Anaerobic | 0.5426149 | 0.6289030 | 0.79 |
| Facultatively Anaerobic | 0.1849250 | 0.3100509 | 0.05 (Borderline significant) |
| Gram Negative | 0.4829289 | 0.3874051 | 1 |
| Gram Positive | 0.5170711 | 0.6125949 | 1 |
| Forms Biofilms | 0.23252654 | 0.08949628 | 0.329004 |
| Contains Mobile elements | 0.7316442 | 0.6303096 | 0.428571 |
| Potentially Pathogenic | 0.3116034 | 0.3926607 | 0.93 |
| Stress-tolerant | 0.0058354205 | 0.0005494533 | 0.14 |

## References

1. Hirai, A.H., et al., *Neonatal Abstinence Syndrome and Maternal Opioid-Related Diagnoses in the US*, *2010-2017*. Jama, 2021. 325(2): p. 146–155.

2. Ko, J.Y., et al., *Vital Signs: Prescription Opioid Pain Reliever Use During Pregnancy – 34 U.S. Jurisdictions*, *2019*. MMWR Morb Mortal Wkly Rep, 2020. 69(28): p. 897–903.

3. Jilani, S.M., et al., *Evaluation of State-Led Surveillance of Neonatal Abstinence Syndrome — Six U.S. States*, *2018–2021*. MMWR Morbidity and Mortality Weekly Report, 2022. 71(2): p. 37–42.

4. Mactier, H. and R. Hamilton, Prenatal opioid exposure - Increasing evidence of harm. Early Hum Dev, 2020. 150: p. 105188.

5. Vassoler, F.M. and M.E. Wimmer, Consequences of Parental Opioid Exposure on Neurophysiology, Behavior, and Health in the Next Generations. Cold Spring Harb Perspect Med, 2021. 11(10).

6. Odegaard, K.E., et al., Characterization of the intergenerational impact of in utero and postnatal oxycodone exposure. Translational Psychiatry, 2020. 10(1): p. 329.

7. Ferrante, J.R. and J.A. Blendy, Advances in animal models of prenatal opioid exposure. Trends in Neurosciences, 2024. 47(5): p. 367–382.

8. Byrnes, E.M. and F.M. Vassoler, Modeling prenatal opioid exposure in animals: Current findings and future directions. Front Neuroendocrinol, 2018. 51: p. 1–13.

9. Fong, J., et al., Developmental Outcomes after Opioid Exposure in the Fetus and Neonate. NeoReviews, 2024. 25(6): p. e325–e337.

10. Jantzie, L.L., et al., Prenatal opioid exposure: The next neonatal neuroinflammatory disease. Brain Behav Immun, 2020. 84: p. 45–58.

11. Newville, J., et al., Perinatal Opioid Exposure Primes the Peripheral Immune System Toward Hyperreactivity. Frontiers in Pediatrics, 2020. Volume 8 - 2020.

12. Wang, F., et al., Morphine induces changes in the gut microbiome and metabolome in a morphine dependence model. Sci Rep, 2018. 8(1): p. 3596.

13. Wang, F. and S. Roy, Gut Homeostasis, Microbial Dysbiosis, and Opioids. Toxicol Pathol, 2017. 45(1): p. 150–156.

14. Zhang, L., et al., Morphine tolerance is attenuated in germfree mice and reversed by probiotics, implicating the role of gut microbiome. Proc Natl Acad Sci U S A, 2019. 116(27): p. 13523–13532.

15. Meng, J., et al., Morphine induces bacterial translocation in mice by compromising intestinal barrier function in a TLR-dependent manner. PLoS One, 2013. 8(1): p. e54040.

16. Banerjee, S., et al., Opioid-induced gut microbial disruption and bile dysregulation leads to gut barrier compromise and sustained systemic inflammation. Mucosal Immunol, 2016. 9(6): p. 1418–1428.

17. Torres, J., et al., Infants born to mothers with IBD present with altered gut microbiome that transfers abnormalities of the adaptive immune system to germ-free mice. Gut, 2020. 69(1): p. 42–51.

18. Maguire, D. and M. Gröer, Neonatal abstinence syndrome and the gastrointestinal tract. Med Hypotheses, 2016. 97: p. 11–15.

19. Raffaeli, G., et al., Neonatal Abstinence Syndrome: Update on Diagnostic and Therapeutic Strategies. Pharmacotherapy, 2017. 37(7): p. 814–823.

20. Piccotti, L., et al., Neonatal Opioid Withdrawal Syndrome: A Developmental Care Approach. Neonatal Netw, 2019. 38(3): p. 160–169.

21. Abu, Y.F., et al., Opioid-induced dysbiosis of maternal gut microbiota during gestation alters offspring gut microbiota and pain sensitivity. Gut Microbes, 2024. 16(1): p. 2292224.

22. Collado, M.C., et al., Human gut colonisation may be initiated in utero by distinct microbial communities in the placenta and amniotic fluid. Sci Rep, 2016. 6: p. 23129.

23. Tanaka, M. and J. Nakayama, Development of the gut microbiota in infancy and its impact on health in later life. Allergol Int, 2017. 66(4): p. 515–522.

24. Wachman, E.M. and L.A. Farrer, The genetics and epigenetics of Neonatal Abstinence Syndrome. Semin Fetal Neonatal Med, 2019. 24(2): p. 105–110.

25. Lu, X., et al., Maternal gut microbiota in the health of mothers and offspring: from the perspective of immunology. Front Immunol, 2024. 15: p. 1362784.

26. Grecco, G.G., et al., Prenatal opioid administration induces shared alterations to the maternal and offspring gut microbiome: A preliminary analysis. Drug Alcohol Depend, 2021. 227: p. 108914.

27. Abu, Y. and S. Roy, Intestinal dysbiosis during pregnancy and microbiota-associated impairments in offspring. Frontiers in Microbiomes, 2025. **Volume** 4 - 2025.

28. Kolli, U. and S. Roy, The role of the gut microbiome and microbial metabolism in mediating opioid-induced changes in the epigenome. Frontiers in Microbiology, 2023. **Volume** 14 - 2023.

29. Jalodia, R., et al., Morphine mediated neutrophil infiltration in intestinal tissue play essential role in histological damage and microbial dysbiosis. Gut Microbes, 2022. 14(1): p. 2143225.

30. Woo, V. and T. Alenghat, Epigenetic regulation by gut microbiota. Gut Microbes, 2022. 14(1): p. 2022407.

31. Brace, L.E., et al., Increased oxidative phosphorylation in response to acute and chronic DNA damage. npj Aging and Mechanisms of Disease, 2016. 2(1): p. 16022.

32. Lee, I. and M. Hüttemann, Energy crisis: The role of oxidative phosphorylation in acute inflammation and sepsis. Biochimica et Biophysica Acta (BBA) - Molecular Basis of Disease, 2014. 1842(9): p. 1579–1586.

33. Reymond, S., et al., Morphine-induced modulation of Nrf2-antioxidant response element signaling pathway in primary human brain microvascular endothelial cells. Scientific Reports, 2022. 12(1): p. 4588.

34. Zahmatkesh, P.M., et al., Impact of opioids on oxidative status and related signaling pathways: An integrated view. Journal of Opioid Management, 2017. 13(4): p. 241–251.

35. Gordon, S., Alternative activation of macrophages. Nature Reviews Immunology, 2003. 3(1): p. 23–35.

36. Leopold Wager, C.M. and F.L. Wormley, Jr., Classical versus alternative macrophage activation: the Ying and the Yang in host defense against pulmonary fungal infections. Mucosal Immunol, 2014. 7(5): p. 1023–35.

37. Gorman, G.S., et al., Mitochondrial diseases. Nat Rev Dis Primers, 2016. 2: p. 16080.

38. Garrido-Pérez, N., et al., Oxidative Phosphorylation Dysfunction Modifies the Cell Secretome. Int J Mol Sci, 2020. 21(9).

39. Davanipour, Z., et al., Endogenous melatonin and oxidatively damaged guanine in DNA. BMC Endocr Disord, 2009. 9: p. 22.

40. Smith, C.J., et al., Microbial modulation via cross-fostering prevents the effects of pervasive environmental stressors on microglia and social behavior, but not the dopamine system. Mol Psychiatry, 2023. 28(6): p. 2549–2562.

41. Giaimo, B.D. and T. Borggrefe, Enhancer-promoter communication: unraveling enhancer strength and positioning within a given topologically associating domain (TAD). Signal Transduction and Targeted Therapy, 2022. 7(1): p. 281.

42. Liu, S., et al., LOF variants identifying candidate genes of laterality defects patients with congenital heart disease. PLOS Genetics, 2022. 18(12): p. e1010530.

43. Yang, J., et al., VSTM2L protects prostate cancer cells against ferroptosis via inhibiting VDAC1 oligomerization and maintaining mitochondria homeostasis. Nature Communications, 2025. 16(1): p. 1160.

44. Yu, L., et al., CD69 enhances immunosuppressive function of regulatory T-cells and attenuates colitis by prompting IL-10 production. Cell Death & Disease, 2018. 9(9): p. 905.

45. Westin, M.A., S.E. Alexson, and M.C. Hunt, Molecular cloning and characterization of two mouse peroxisome proliferator-activated receptor alpha (PPARalpha)-regulated peroxisomal acyl-CoA thioesterases. J Biol Chem, 2004. 279(21): p. 21841–8.

46. Giovannini, E., et al., Fetal and Infant Effects of Maternal Opioid Use during Pregnancy: A Literature Review including Clinical, Toxicological, Pharmacogenomic, and Epigenetic Aspects for Forensic Evaluation. Children, 2024. 11(3): p. 278.

47. Borrelli, K.N., et al., Effect of Prenatal Opioid Exposure on the Human Placental Methylome. Biomedicines, 2022. 10(5).

48. Radhakrishna, U., et al., Placental DNA methylation profiles in opioid-exposed pregnancies and associations with the neonatal opioid withdrawal syndrome. Genomics, 2021. 113(3): p. 1127–1135.

49. https://www.alzet.com/iprecio-pump/iprecio-overview/, 2025.

50. Abu, Y., et al., Brief Hydromorphone Exposure During Pregnancy Sufficient to Induce Maternal and Neonatal Microbial Dysbiosis. J Neuroimmune Pharmacol, 2022. 17(1-2): p. 367–375.

51. Lyu, Z., et al., Long-Term Effects of Developmental Exposure to Oxycodone on Gut Microbiota and Relationship to Adult Behaviors and Metabolism. mSystems, 2022. 7(4): p. e0033622.

52. Antoine, D., et al., Neonatal Morphine Results in Long-Lasting Alterations to the Gut Microbiome in Adolescence and Adulthood in a Murine Model. Pharmaceutics, 2022. 14(9).

53. Fusco, W., et al., Short-Chain Fatty-Acid-Producing Bacteria: Key Components of the Human Gut Microbiota. Nutrients, 2023. 15(9).

54. O’Callaghan, A. and D. van Sinderen, Bifidobacteria and Their Role as Members of the Human Gut Microbiota. Frontiers in Microbiology, 2016. **Volume** 7 - 2016.

55. Hanski, E., A. Raulo, and S.C.L. Knowles, Early-life gut microbiota assembly patterns are conserved between laboratory and wild mice. Communications Biology, 2024. 7(1): p. 1456.

56. Huang, Y.E., et al., Disrupted establishment of anaerobe and facultative anaerobe balance in preterm infants with extrauterine growth restriction. Front Pediatr, 2022. 10: p. 935458.

57. Friedman, E.S., et al., Microbes vs. chemistry in the origin of the anaerobic gut lumen. Proc Natl Acad Sci U S A, 2018. 115(16): p. 4170–4175.

58. Daft, J.G., et al., Cross-fostering immediately after birth induces a permanent microbiota shift that is shaped by the nursing mother. Microbiome, 2015. 3: p. 17.

59. Sikder, M.A.A., et al., Maternal diet modulates the infant microbiome and intestinal Flt3L necessary for dendritic cell development and immunity to respiratory infection. Immunity, 2023. 56(5): p. 1098–1114.e10.

60. Alsulaiman, R.M., et al., Gut microbiota analyses of inflammatory bowel diseases from a representative Saudi population. BMC Gastroenterol, 2023. 23(1): p. 258.

61. Dai, W., et al., Uncovering a causal connection between the Lachnoclostridium genus in fecal microbiota and non-alcoholic fatty liver disease: a two-sample Mendelian randomization analysis. Frontiers in Microbiology, 2023. **Volume** 14 **-** 2023.

62. Zhu, A., et al., Diagnosis and functional prediction of microbial markers in tumor tissues of sporadic colorectal cancer patients associated with the MLH1 protein phenotype. Frontiers in Oncology, 2023. **Volume** 12 **-** 2022.

63. Zhang, H., et al., Targeting CDK9 Reactivates Epigenetically Silenced Genes in Cancer. Cell, 2018. 175(5): p. 1244–1258.e26.

64. Buenrostro, J.D., et al., ATAC-seq: A Method for Assaying Chromatin Accessibility Genome-Wide. Curr Protoc Mol Biol, 2015. 109: p. 21.29.1–21.29.9.

65. Smith, R.J., et al., Single-cell chromatin profiling of the primitive gut tube reveals regulatory dynamics underlying lineage fate decisions. Nature Communications, 2022. 13(1): p. 2965.

66. Liu, S.X., et al., Differential gene expression and chromatin accessibility in the medial prefrontal cortex associated with individual differences in rat behavioral models of opioid use disorder. bioRxiv, 2024.

67. Zhra, M., et al., Sirtuins and Gut Microbiota: Dynamics in Health and a Journey from Metabolic Dysfunction to Hepatocellular Carcinoma. Cells, 2025. 14(6).

68. Shalapour, S., et al., Interleukin-7 Links T Lymphocyte and Intestinal Epithelial Cell Homeostasis. PLOS ONE, 2012. 7(2): p. e31939.

69. Meduri, G.U., Synergistic glucocorticoids, vitamins, and microbiome strategies for gut protection in critical illness. Exploration of Endocrine and Metabolic Diseases, 2025. 2: p. 101432.

70. Botía-Sánchez, M., M.E. Alarcón-Riquelme, and G. Galicia, B Cells and Microbiota in Autoimmunity. Int J Mol Sci, 2021. 22(9).

71. Elkhalil, A., A. Whited, and P. Ghose, SQST-1/p62-regulated SKN-1/Nrf mediates a phagocytic stress response via transcriptional activation of lyst-1/LYST. PLOS Genetics, 2025. 21(5): p. e1011696.

